# JOnTADS: a unified caller for TADs and stripes in Hi-C data

**DOI:** 10.1101/2024.11.06.622323

**Authors:** Qiuhai Zeng, Guanjue Xiang, Yu Zhang, Ross C. Hardison, Qunhua Li

## Abstract

Topologically associating domains (TADs) and stripes are important architectural structures on Hi-C data that are important for gene regulation. We present Joint Optimized nested TADs and Stripes (JOnTADS), a unified caller for TADs and stripes in Hi-C data. JOnTADS effectively identifies hierarchical TADs and stripes in population Hi-C and micro-C datasets, and hierarchical TADs in single-cell Hi-C data. It provides robust identifications aligned with known biology and effectively captures interaction frequency variations in contact maps across diverse contexts. When multiple samples are available, JOnTADS leverages shared information across samples in TAD identification, reducing unwanted variation in TAD boundary identification while preserving biological differences. This approach enables robust identifications in single-cell Hi-C data, effectively addressing challenges posed by data sparsity. JOnTADS is computationally efficient and requires minimal user tuning.

## Introduction

The three-dimensional (3D) organization of the genome plays a crucial role in gene regulation. Chromosome conformation capture techniques, notably Hi-C, unveil the hierarchical folding of chromatin across multiple scales. At the gene locus level, i.e., hundreds of kilo bases, chromatin organizes into distinct high-interaction regions, known as topologically associating domains (TADs), representing local regions where genomic loci interact more frequently [1]. TADs are known to harbor cell-type specific frequently interacting region [2] and are correlated with varying transcriptional activity [3], suggesting that TADs have a potential role in gene regulation by facilitating enhancer–promoter interaction and insulating regulatory activities [4]. Disruptions in TAD structures have been linked to various diseases [5, 6, 7, 8], although some disruptions exhibit only mild effects on gene expression [9, 10, 11, 12]. The precise functional contribution of TADs to gene regulation remains an open question [4]. Another 3D structure at the gene locus level, known as architectural stripes, has been observed to be associated with enhancer activity and gene activation [13, 14]. These stripes manifest as asymmetric patterns of contact spanning a contiguous genomic interval over several hundred kilobases in Hi-C maps [13]. They have been shown to link active enhancers near a strong boundary to gene activation [13]. Given their potential impact on cell identity and function, accurate detection of these features from Hi-C and other chromatin conformation measurements is crucial.

In recent years, a large volume of chromosome conformation data has been produced [15, 16], including Hi-C data obtained from diverse cell types and conditions. This wealth of data allows for the comparison of domain structures across different conditions and the exploration of TAD dynamics’ impact on cellular functions [17, 18, 15]. Advances in high resolution genome-wide pairwise chromatin conformation technologies, such as Micro-C, reveal intricate stripes and other interaction patterns within TADs [19, 20], providing new insights into the role of TADs internal structures and stripes in gene regulation. Additionally, single-cell genome-wide interaction technologies [21, 22, 23] enable the examination of cell-specific TAD-like domain (TLD) structures at a single-cell resolution [24, 25], offering the potential to uncover connections between genome structure and function in diverse cellular environments [26].

Despite the abundance of available data resources, robustly identifying architectural structures from diverse Hi-C data and conducting comparative analyses across samples remain computational challenges. Presently, TAD identification and stripe identification are regarded as separate problems, requiring distinct software for each task.

Various algorithms have been developed for TAD identification from Hi-C data [27, 28, 29, 30, 31, 32, 33, 34] (See [35, 36, 37, 38] for a comprehensive review and comparison). However, these algorithms are mainly tailored for a single contact matrix, with the majority optimized for low-resolution population Hi-C data. Challenges arise when applying these algorithms to compare TAD structures across multiple samples. Low reproducibility has been observed across the TAD boundaries identified on replicate samples [39, 40], partially due to many small shifts between TAD boundaries across samples (see an example in Supplementary Fig. 1a). This is likely due to the callers’ lack of robustness against stochastic variation in the data, hampering the accurate identification of biologically significant alterations in chromatin organization across samples [41]. Additionally, different data types possess unique characteristics, making algorithms developed for population Hi-C less effective for sparse or high-resolution data. For instance, single-cell Hi-C data, characterized by ultra sparsity and a lower signal-to-noise ratio, poses challenges in distinguishing meaningful chromatin organization differences from technical noise. Currently, only deTOKI [34] and deDoc2 [42] are specifically tailored for single-cell Hi-C data, of which, deTOKI only provides boundary positions for TAD-like structures but not their hierarchy. Similarly, Micro-C, with considerably larger data sizes due to its high resolution, demands greater computational efficiency and memory usage from the identification algorithm.

**Figure 1:**
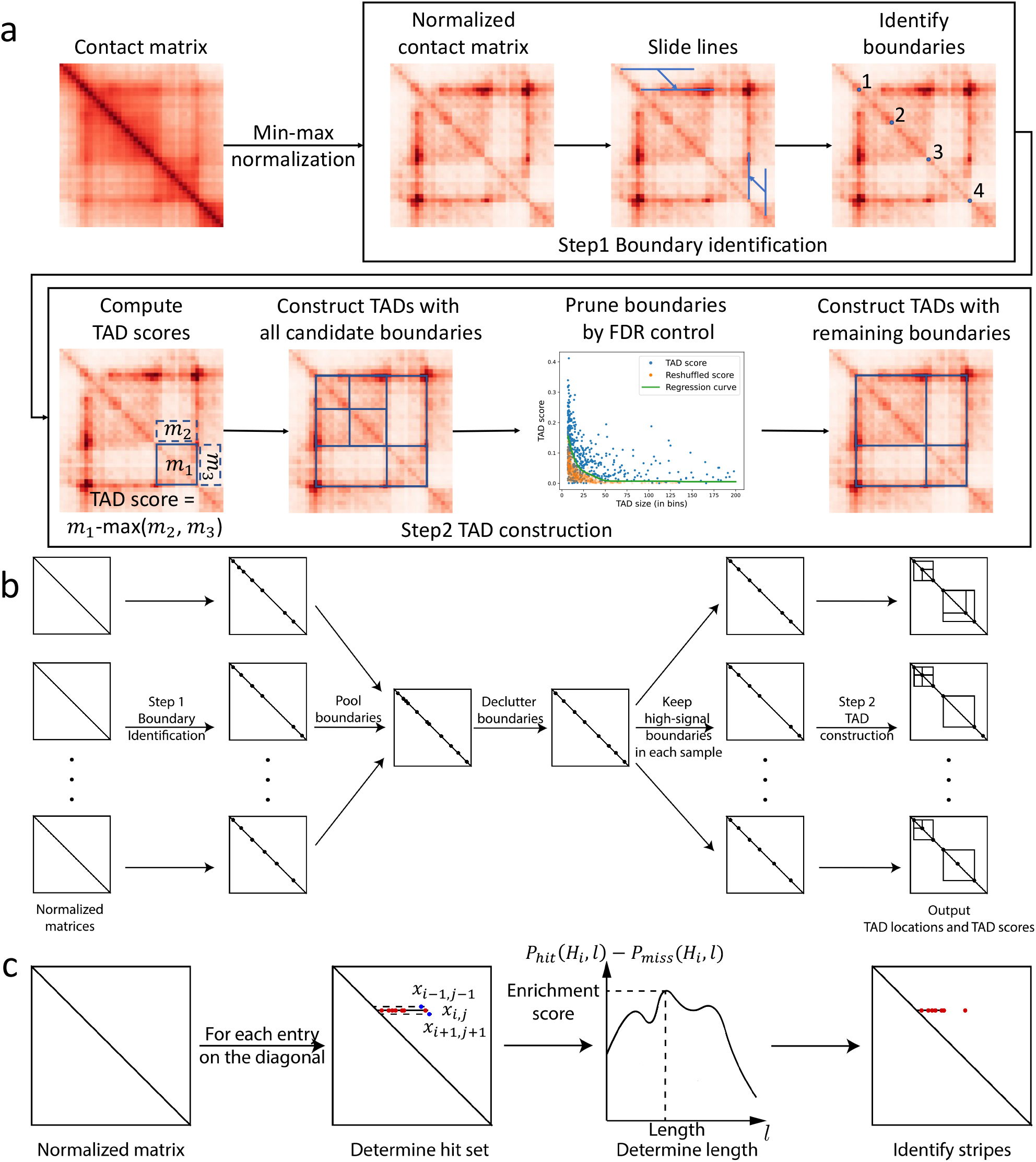
Overview of JOnTADS’s TAD and stripe calling procedure. **a** TAD calling on a single contact matrix. **b** Joint TAD calling across multiple samples. **c** Stripe calling procedure.

Existing computational methods for stripe identification also come with limitations. For example, Zebra lacks a published model implementation [13], DomainClassifyR detects stripe domains solely at TAD boundaries [14], and the recent method, Stripenn, is confined to data below 5Kb resolution [43]. Therefore, there remains a need for accurate and efficient architectural stripe detection across various cell types and conditions. Moreover, the requirement for separate software for identifying TADs and stripes can be inconvenient for users. Researchers often desire to study the roles for both structures in gene regulation, and using different software tools for their identification may introduce complications due to distinct preprocessing procedures and output formats across software platforms.

In this study, we develop the **J**oint **O**ptimized **n**ested **TAD** and **S**tripe (JOnTADS) model, a unified TAD and stripe caller that addresses the aforementioned challenges across diverse genome-wide chromatin conformation data, such as population Hi-C, single-cell Hi-C and micro-C data. JOnTADS is the first method capable of performing joint TADs calling across multiple contact matrices, leveraging information across distinct samples to mitigate undesired shifts in TAD boundary locations and thereby accentuating biologically relevant variations. We demonstrate the effectiveness of JOnTADS across a spectrum of data types, spanning single contact maps, multiple contact maps from different cell types and cell states, and single cell and high-resolution Micro-C data.

## Results

### Overview of the JOnTADS algorithm

JOnTADS is a novel method for identifying hierarchical TADs and stripes in various chromatin conformation capture data, including population Hi-C, single-cell Hi-C and micro-C. It is applicable in both standard analyses of a single contact matrix and more complex scenarios, such as comparing and joint analyzing contact maps across multiple conditions, time points, or cell types, including single-cell Hi-C data.

JOnTADS takes one or multiple raw count contact matrices as input, and uses a modified min-max normalization to standardize each contact matrix. This ensures comparability of signals across different genomic distances and matrices. The normalized matrices then serve as the basis for identifying TADs and stripes.

JOnTADS identifies TADs on a single contact matrix through a two-step process (Figure 1a). In the first step, it scans the contact map along its diagonal using lines of varying lengths to detect potential boundaries. These are identified at locations where there is a strong contrast between signals on the line and its neighbors. Unlike the typical square-shaped scanning window that captures regions with large differences in average signals against their surroundings, JOnTADS’s line-shaped scan pinpoints TAD edges. This approach is particularly effective at identifying TAD structures with weaker internal interactions compared to those at the edges, a pattern commonly seen in sparse datasets like single-cell Hi-C (Supplementary Fig. 1b).

In the second step, JOnTADS assembles these candidate boundaries into TADs by optimizing the fit between the estimated TAD hierarchy and observed interaction pattern using a dynamic programming algorithm. It then prunes TADs without significant contact frequency enrichment by estimating the relationship between mean contact frequency and TAD size using a shuffled contact matrix and applying size-specific thresholds to filter out putative identifications at a given false discovery rate. This TAD-size-adaptive pruning ensures the method’s flexibility, particularly for sparse or high-resolution Hi-C data, which show a distinct correlation between TAD size and mean contact frequency that is not present in population Hi-C data (Supplementary Fig. 2).

**Figure 2:**
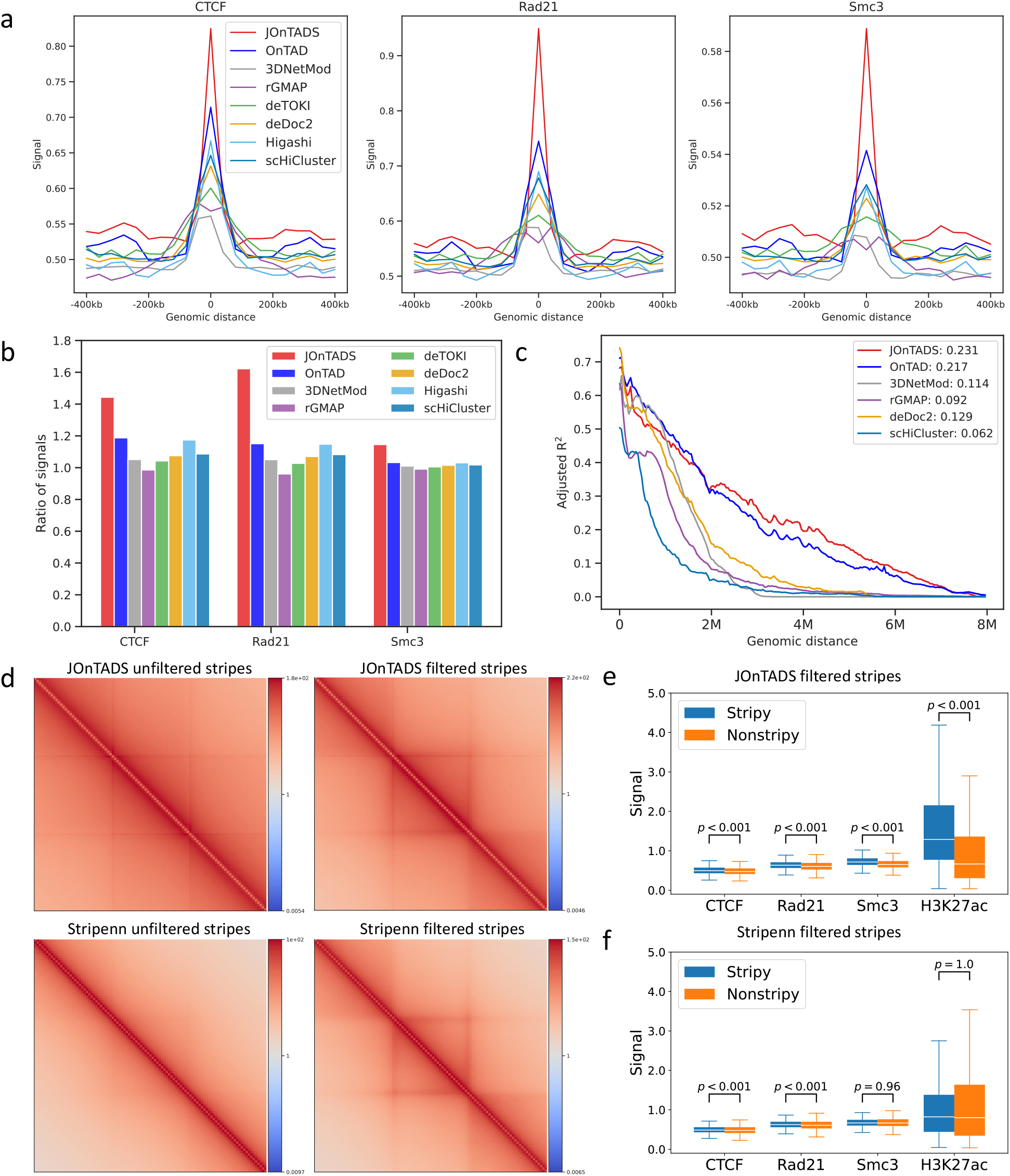
Comparison of TAD and stripe identification in the bulk-cell GM12878 Hi-C data. **a** Average CTCF, Rad21, and Smc3 ChIP-seq signal at identified TAD boundaries and surrounding ±10 bins. **b** Ratio of CTCF, Rad21, and Smc3 signals between TAD boundaries and their adjacent ±1 bins. **c** TAD adj-*R*^2^ at different genomic distances. **d** Pileup plots of stripes detected by JOnTADS and Stripenn. Filtered: stripes passing the filter of Stripenn p-values ≤0.1. Unfiltered: all identified stripes. **e-f** Distributions of average CTCF, Rad21 and Smc3 signals within stripy and non-stripy TADs based on filtered stripes identified by JOnTADS (**e**) or Stripenn (**f**). The p-values were calculated using one-sided t-test.

When multiple contact matrices are available, JOnTADS first identifies candidate boundaries from each matrix as described in Step 1 above, and then performs a joint TAD calling procedure that integrates information across matrices (Figure 1b). The procedure consolidates boundaries at similar locations across samples using a dynamic programming algorithm. High-signal boundaries in individual samples that align with these consolidated boundaries are then used to construct TADs, as outlined in Step 2, allowing for distinct TADs to be called in different samples while ensuring more robust boundary identification.

To identify potential stripes, JOnTADS first conducts a line scan along the matrix diagonal, detecting stripes by analyzing contrasts in signal strength along the line relative to neighboring regions (Figure 1c). It then uses a novel scoring approach, inspired by gene set enrichment analysis (GSEA) [44], to determine the endpoint and the strength of a stripe. Specifically, JOnTADS tracks the cumulative proportion of bins with and without enriched signals along a stripe, moving outwards from its starting point. The maximum disparity between these proportions, signaling a reduction of signal enrichment, defines the stripe’s endpoint. Each candidate stripe is then assigned an enrichment score that reflects its signal strength relative to adjacent regions.

JOnTADS computes the cumulative fractions of bins with and without signal enrichment over adjacent regions along a candidate stripe, moving outward from the diagonal. The maximum disparity in these cumulative fractions, marking the reduction of signal enrichment, is used to define the stripe’s endpoint. Each candidate stripe is then assigned an enrichment score that reflects its signal strength relative to adjacent regions.

JOnTADS is computationally efficient and requires minimal user input (Supplementary ‘Parameter choice’), adapting to various contexts automatically without the need for users to tune parameters due to its data-driven procedures. The running time comparison in Supplementary ‘Time complexity’ and Supplementary Table 1 demonstrates its efficiency, especially for high-resolution data.

### TAD and stripe identification on a single population Hi-C dataset

We first assessed the performance of JOnTADS on a single population Hi-C dataset at a resolution commonly targeted by most existing algorithms, using the well-studied GM12878 dataset [28]. We compared TAD calling with OnTAD, rGMAP, 3DNetMod and DeTOKI at 40kb resolution and compared stripe calling at 5kb resolution with Stripenn [43], a recently published stripe caller known for its superior performance over other existing stripe callers. Among these, rGMAP identified the fewest TADs (around 2,500), OnTAD and JOnTADS identify around 4,500-6,000 TADs, while 3DNetMod identified over 15,000 TADs, significantly more than the other methods (Supplementary Table 2).

To evaluate the accuracy of TAD boundary identification, we calculated the average signal of CTCF and two cohesin complex proteins, Rad21 and Smc3, at boundary positions and their surrounding ±10 bins. JOnTADS showed the highest signals at boundary positions for all three proteins (Figure 2a) and also the highest contrast in average boundary protein signals compared to adjacent (±1) bins (Figure 2b), affirming its precision in boundary identification.

To assess the accuracy of TAD hierarchy construction, we used TAD-adjR^2^ [33] to measure how well the variation of Hi-C signals in a contact matrix can be explained by the TADs (subTADs) and non-TAD classification assigned by each caller. JOnTADS achieved the highest overall TAD-adjR^2^ (0.231), substantially surpassing OnTAD (0.217), rGMAP (0.084), and 3DNetMod (0.114) (Figure 2c), demonstrating that JOnTADS effectively identified biologically meaningful domain structures in a single population Hi-C sample.

Furthermore, the stripes identified by JOnTADS showed a higher overall contrast compared to their adjacent regions than those identified by Stripenn, as seen in the pileup plots (Figure 2d) [45], indicating stronger average interaction intensity within the identified stripes. After applying Stripenn’s p-value filter (*<* 0.1), which assessed the significance of the contrast between a stripe and its neighboring bins, the filtered stripes from both methods exhibited similar levels of enrichment relative to their surrounding regions.

To confirm biological relevance of the stripes identified by JOnTADS, we investigated the signal of architectural proteins and the active enhancer mark, H3K27ac, in TAD containing stripes. Previous studies have shown that CTCF and cohesin subunits are highly enriched at stripe anchors [14, 13], and that TADs harboring stripes tend to have a higher signal of these architectural proteins than TADs without stripes in activated B cells and a higher level of H3K27ac in double positive thymocytes [43]. We classified TADs as stripy if they contained at least one stripe within their region and as non-stripy if they did not. We found that when stripy TADs were defined using stripes identified by JOnTADS, stripy TADs showed significantly higher average signal of CTCF, Rad21, Smc3 and H3K27ac than non-stripy TADs (Figure 2e-f). However, for TADs classified using stripes identified by Stripenn, only CTCF and Rad21 signals showed significant signal in stripy TADs, while no significant signal was observed for Smc3 and H3K27ac. These results confirmed the precision and biological relevance of JOnTADS’s stripe identifications.

### Comparing TAD hierarchies across multiple cell types in a cell lineage

Next, we evaluated JOnTADS’s performance on multiple samples by assessing its joint TAD calling accuracy using a bulk human ES cell (hESC) lineage dataset from [46]. This lineage comprises human embryonic stem (ES) cells and four derived cell types: mesendoderm (ME), mesenchymal stem cells (MS), neural progenitor cells (NP), and trophoblast-like cells (TB), with two biological replicates for each cell type. As differentiation progresses, the similarity between ES and its derived cell type is expected to decrease with the differentiation times, shown as follows [47],

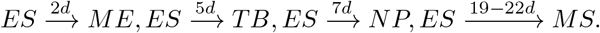

This distance in lineage relationship is expected to be reflected in both the similarity between contact matrices and the similarity between TADs structures. We used JOnTADS to jointly call TADs across all ten contact matrices (Methods) and applied other methods to each contact matrix individually. Hierarchical clustering was performed for each method to infer interrelationships between cell types, based on the pairwise Jaccard Index for TADs for population data (Methods), which measured the similarity of identified TAD hierarchies between sample pairs.

While all methods correctly grouped replicates together, only JOnTADS and 3DNetMod produced TAD hierarchies that aligned with the lineage relationship indicated by differentiation time (Figure 3a). Additionally, JOnTADS was the only method that consistently demonstrated a higher correlation between replicate samples compared to non-replicate samples. Notably, JOnTADS consistently exhibited considerably higher Jaccard Index values than all other methods across all sample pairs, likely due to its ability in minimizing boundary shifting across samples.

**Figure 3:**
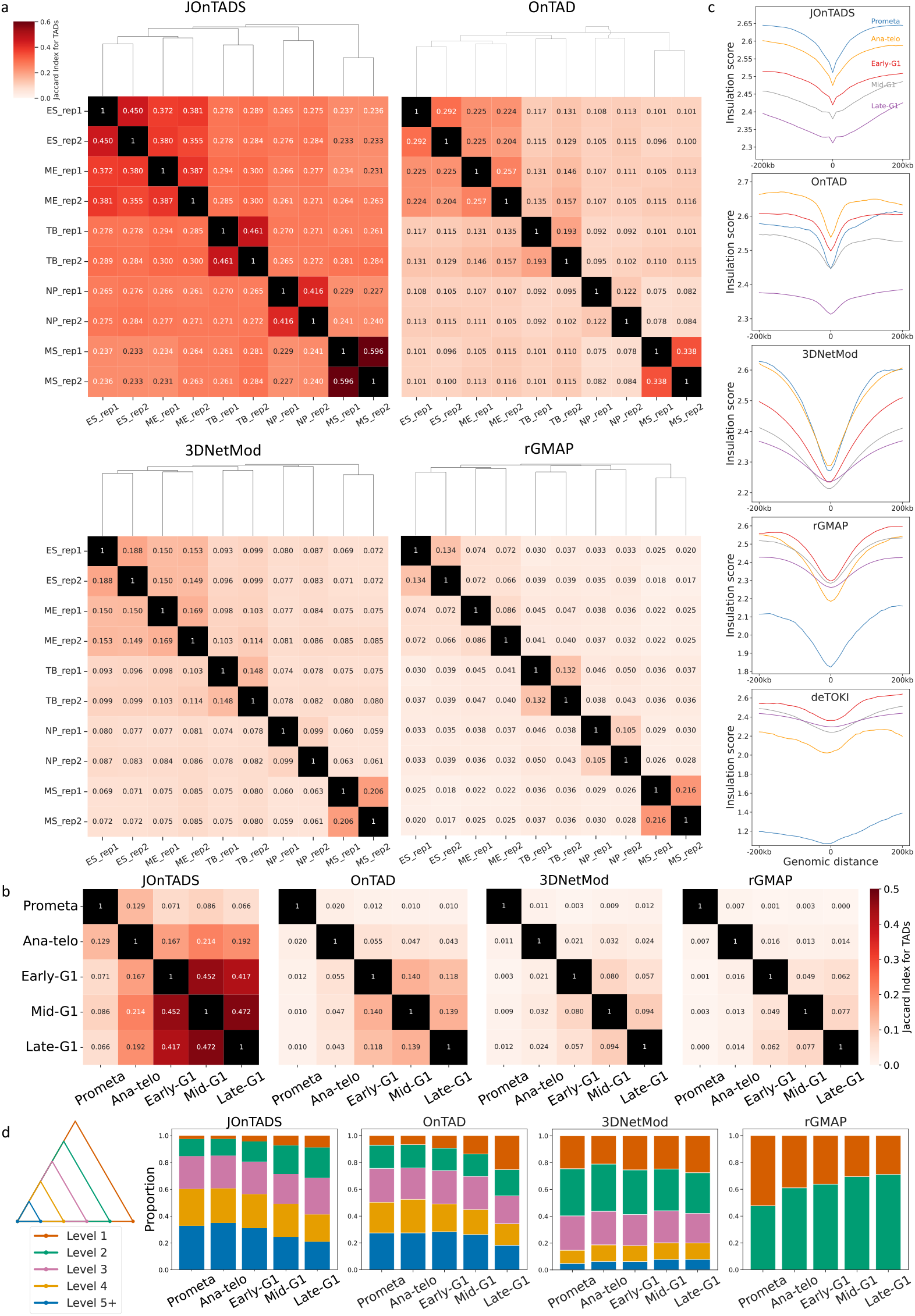
Comparison of TAD identification in two multi-sample bulk-cell Hi-C datasets. **a** Jaccard Index for TADs (JIT) between each pair of samples in the bulk-cell hESC lineage data. **b-c** JIT in bulk-cell G1E-ER4 cell cycle data and insulation scores of newly emerged boundaries. **b** Jaccard Index between each pair of phases. **c** Average insulation scores of newly emerged boundaries and their adjacent regions (±20 bins) at each phase. Insulation scores are calculated using late G1 phase data with window size of 100. **d** Proportion of newly emerged boundaries at different TAD levels at each phase. In **c** and **d**, only the newly emerged boundaries that persist through the rest of the phases are included.

### Analysis of TAD formation dynamics in different phases of cell cycles

We next used JOnTADS to analyze dynamic changes in chromosome reorganization during the transition from mitosis to G1 in mouse erythroid cell populations [18]. This dataset includes highly purified, synchronous mouse G1E-ER4 cells in five cell cycle phases: two phases in mitosis (prometaphase and anatelophase) and three G1 phases (early-G1, mid-G1, and late-G1). During prometaphase, chromatin within a cell is in a relatively unorganized state with a substantial loss of domain structures. As the cell cycle progresses into late G1, contact domains gradually form into a fully established chromatin organization. It has been reported [18] that the hierarchical domain reorganization during the transition from prometaphase to G1 follows a ‘bottom-up’ model, where nested domain structures emerge from the convergence of subTADs formed in previous stages. As in the previous section, we ran JOnTADS on all five phases jointly and applied other methods to each contact matrix separately. For 3DNetMod, we used the published TAD calling results from [18]. We evaluated the similarity of TAD hierarchies between each pair of phases using the Jaccard Index for TADs for population data (Methods).

As shown in Figure 3b, the Jaccard Indices within the three G1 stages (0.41-0.47) were substantially higher than those between mitosis and G1 stages (0.07-0.21) for TAD hierarchies identified by JOnTADS, suggesting a drastic difference in the chromatin conformation between mitotic and G1 stages. This finding aligned with previous reports that interphase chromatin exhibited a highly compartmentalized organization, while mitotic phases entirely lacked this structure [48, 49]. Other methods also showed a similar trend, but the distinctions were much smaller (within G1: 0.05-0.14; between mitosis and G1: 0.00-0.06). For example, Jaccard Indices from 3DNetMod and rGMAP remained low across all stages, even between the three G1 stages (*<* 0.1), making it difficult to confidently differentiate mitosis and G1 stages. Notably, JOnTADS consistently showed much higher Jaccard Indices than other methods for all pairs of phases, likely due to its ability to minimize unwanted boundary-calling variation using combined information across samples, thereby enhancing the biological distinctions between mitosis and G1 stages.

To evaluate the biological relevance of identified TADs across the cell cycle, we compared the insulation at their boundaries at different stages. According to the bottom-up model, smaller subTADs form in the early stages and gradually merge into larger multi-domain TADs as cells progress from prometaphase to G1 [18]. Consequently, boundaries formed in earlier stages, likely associated with subTADs, tend to exhibit higher insulation scores [29, 50] (indicative of a lower insulation) than those formed in later stages, which likely contribute to TADs encompassing subTADs. Here we focused on the boundaries that persisted through the late-G1 and categorized them based on the stage of initial detection. We then computed their insulation scores and those of surrounding regions (±20 bins) at the late G1 stage. Supplementary Table 3 summarized the number of boundaries in each category and their insulation score ratios relative to adjacent bins.

Indeed, the insulation scores of TAD boundaries identified by JOnTADS and 3DNetMod followed the expected order, with the highest scores for those emerging at prometaphase and the lowest for those emerging at late-G1. In contrast, boundaries identified by OnTAD, rGMAP and DeTOKI did not follow this trend (Figure 3c). Moreover, JOnTADS showed the largest reduction in insulation score at its boundaries relative to adjacent (±1) bins (Supplementary Table 3).

As the cell cycle progresses from mitosis to late-G1, sub-TADs merge into outer TADs [18], likely resulting in an increase in the proportion of newly formed boundaries at the outer TADs. Following [33], we defined TAD levels in an outer-to-inner manner, where the outermost TAD was assigned as level 1, with the levels increasing inward. As expected, both JOnTADS and OnTAD exhibited the trend that the proportion of newly formed boundaries in outer TADs (i.e., lower-level TADs) increased as the cells progressed through different stages of the cell cycle (Figure 3d).

### Analyzing TAD-like domains in Dip-C data

We next evaluated JOnTADS’s ability to identify TAD-like domains (TLD) in single cell chromatin conformation capture data using a Dip-C dataset collected from 14 GM12878 cells [24]. Dip-C data, which captures high-resolution 3D genome structures of single diploid cells, is sparse, containing many zeros and significantly lower read counts compared to population-level data. (See Supplementary Table 4 for a comparison of read counts of this Dip-C data and the GM12878 population Hi-C data in [28]). We compared JOnTADS with OnTAD, rGMAP, and several methods designed for analyzing single-cell Hi-C data, namely deTOKI, deDoc2, Higashi and scHiCluster [34, 42, 26, 51]. As in the previous sections, we ran JOnTADS on the 14 single cell contact matrices jointly and ran other methods on each contact matrix separately. Supplementary Table 2 summarized the numbers of TLDs and TLD boundaries identified by each method.

### JOnTADS excels in identifying biologically relevant TLD boundaries and capturing domain variation across cells

Across all methods, we observed that a large proportion of boundaries were unique to individual cells (21% for deDoc2 and 41 − 67% for all other methods), with less than 2% shared by six of more cells (Figure 4a and Supplementary Table 5), highlighting the high variability of chromatin conformation in single cells.

**Figure 4:**
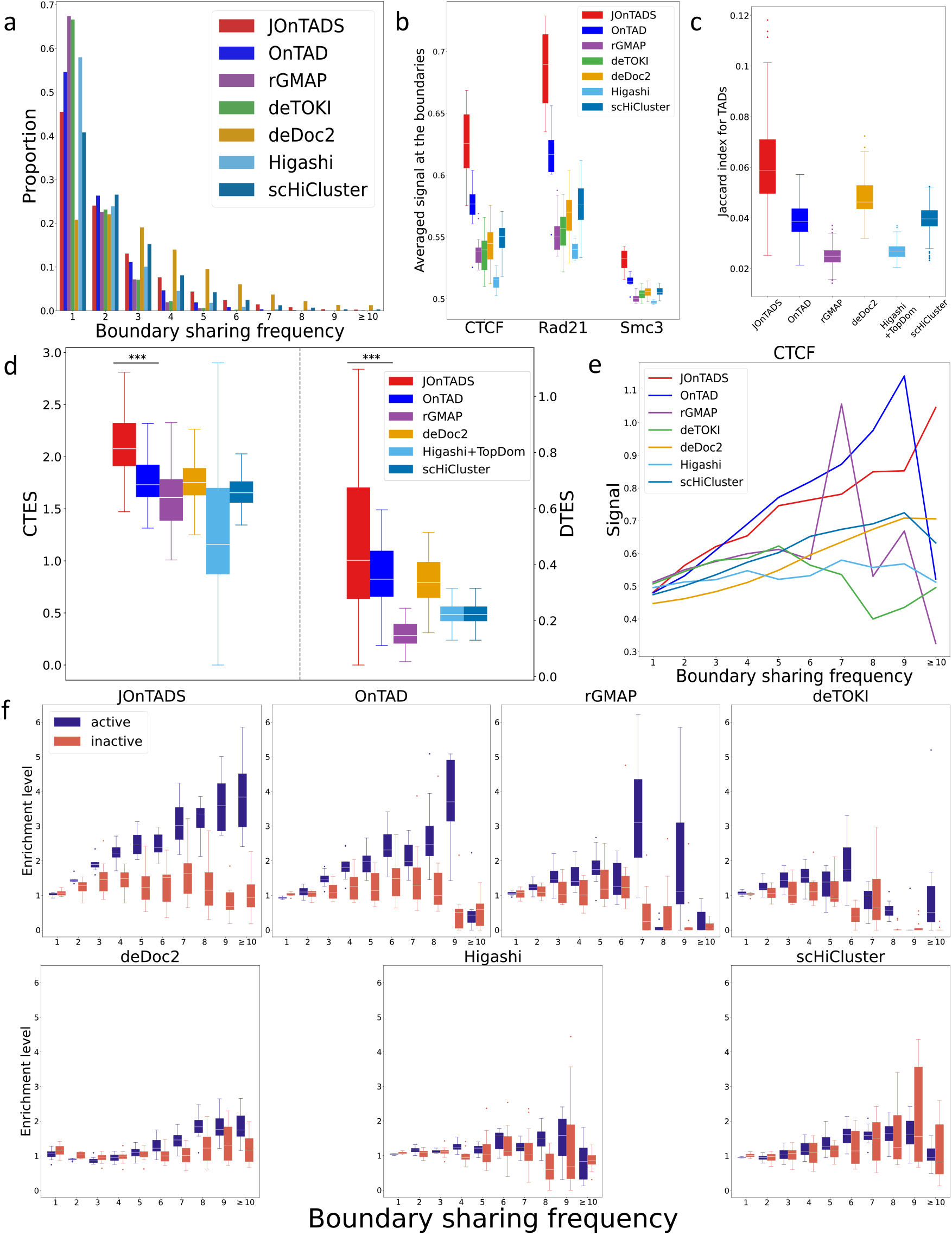
Comparison of TAD identification in the Dip-C GM12878 dataset. **a** Proportion of boundaries with different sharing frequency. **b** Average CTCF, Rad21 and Smc3 signals at single cell boundaries in different single cells. **c** Jaccard Index for TADs (scJIT) between each pair of single cells. **d** CTES and DTES scores between every pair of cells’ TAD results. Higashi+TopDom refers to using Higashi for contact map imputation and using TopDom to identify TADs. ***: *p <* 0.001 obtained from one-sided t-test. **e** CTCF signal at boundaries with different boundary frequencies. **f** Averaged inactive/active IDEAS states signal at boundaries with different boundary frequencies.

To verify the identified TLD boundaries, we examined ChIP-seq signals of boundary proteins (CTCF, Rad21, and Smc3) at the identified TLD boundaries. For all three proteins, JOnTADS consistently showed substantially higher signals than other methods (Figure 4b), confirming its effectiveness in detecting biologically relevant boundaries in single-cell data.

We then examined the similarity of identified domain structures across different single cells. Due to the high variability in chromatin conformation and data sparsity, existing similarity metrics struggle to effectively capture differences in similarity. To address this, we introduced three metrics: a modified Jaccard Index for TADs for single cell data (scJIT), which measured similarity in both domains and boundaries between cell pairs; the Common TLD Enrichment Score (CTES), which quantified the enrichment of contact signals within TLDs shared across cells; and the Differential TLD Enrichment Score (DTES), which measured the contrast in contact signals within differential domain regions between cells (Methods). JOnTADS consistently achieved the highest scJIT, CTES, and DTES among all methods (Figure 4c-d), indicating greater consistency in identified TLD structures, higher signal enrichment in shared TLD regions, and more distinct signal differences in differential TLD regions compared to other methods.

### JOnTADS reveals strong association between boundary sharing and epigenomic characteristics

We investigated whether boundaries shared by more cells exhibited distinct biological characteristics compared to those shared by fewer cells by comparing their CTCF signals and the enrichment of active epigenomic states, which were defined based on the absence of H3K27me3 and H3K9me3 signals [52], derived from combined epigenomic marks from IDEAS [53, 52] (Methods). We found a strong positive correlation between boundary sharing frequency and both CTCF signals and active epigenomic states for boundaries identified by JOnTADS and OnTAD (Figure 4e-f), suggesting a link to CTCF-directed interactions. In contrast, boundaries from other methods lacked this association (Figure 4e-f).

### Analyzing TAD-like domains in single cell Hi-C data

Next we evaluated the performance on single cell Hi-C data from human brain prefrontal cortex [54]. This dataset consists of a total of 4,238 cells, with 14 annotated cell types from four categories, glutamatergic neurons (L23, L4, L5, L6), GABAergic neurons (Ndnf, Vip, Pvalb, Sst), non-neuronal cells (Astro, ODC, OPC) and non-neural cells (MG, MP, Endo). For each cell type, we selected 5 cells with the highest sequencing depth, yielding 70 cells in total. We ran JOnTADS 14 times, once for each cell type, and compared it to OnTAD, rGMAP, deDoc2, Higashi, and scHiCluster [33, 31, 42, 26, 51], which were run on each cell individually. For Higashi, an imputation method that does not call domains, we followed [51] to first use it to impute the contact maps across the 70 cells and used TopDom to identify TLDs for each cell in the imputed maps. We then computed scJIT, CTES, and DTES across all pairs of cells for identified TLDs.

For cells of the same type, JOnTADS achieved the highest scJIT, CTES, and DTES (Figure 5a-b) among all methods, with CTES and DTES significantly surpassing those of other approaches. Notably, JOnTADS showed a much greater distinction in scJIT between cells of the same type and those of different types (Figure 5a), highlighting its capacity to identify TLDs that distinguish cell types. Hierarchical clustering based on scJIT further demonstrated that JOnTADS accurately grouped cells of the same type, separated non-neural from non-neuronal cells, and clustered non-neural types, whereas other methods failed to do so (Figure 5c). Together, these further confirmed JOnTADS’s effectiveness in identifying both shared and differential domain structures and its capacity to identify TLDs that distinguish cell types.

**Figure 5:**
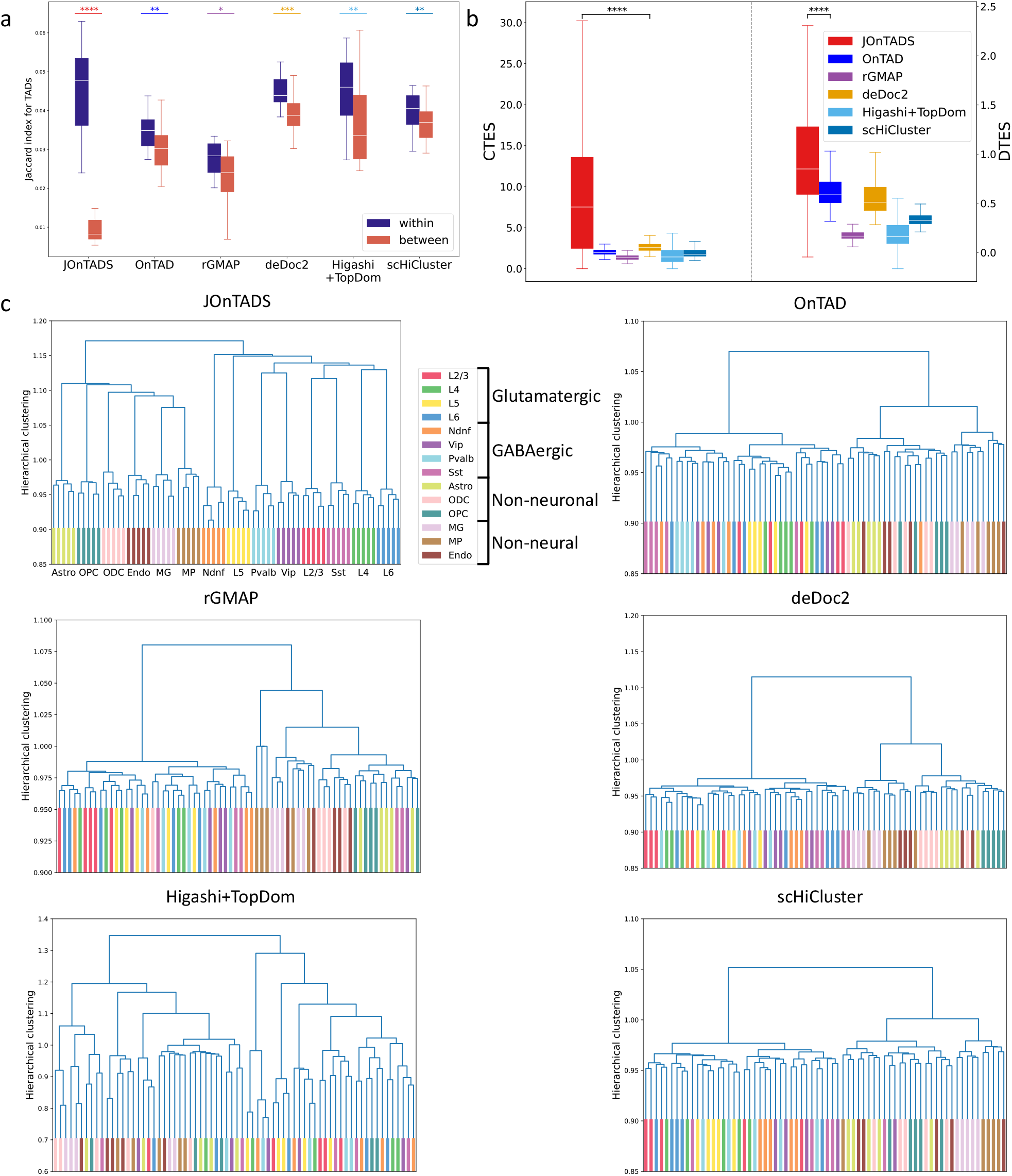
Comparison of TAD identification in the single-cell HiC dataset from human brain prefrontal cortex. **a** Jaccard index for TAD (scJIT) between each pair of single cells. *: *p <* 0.05, **: *p <* 0.01, ***: *p <* 10^−3^, ****: *p <* 10^−6^ **b** CTES and DTES scores between every pair of cells’ TAD results. **c** Hierarchical clustering with the ward method using scJIT as the metric. Higashi+TopDom refers to using Higashi for contact map imputation and using TopDom to identify TADs. The p-values were calculated using one-sided t-test.

### Identifying TADs and stripes on micro-C data

#### JOnTADS captures TAD structures and reveals epigenomic insights from Micro-C contact map

We assessed JOnTADS’s TAD calling performance on the Micro-C H1-hESC dataset with a resolution of 1kb [19]. Because the size of a Micro-C contact matrix can easily exceed tens of gigabytes for a single chromosome, we chose to evaluate JOnTADS on the smallest chromosome (chromosome 22), consisting of 50,819 × 50,819 bins and a size of 4.9 gigabytes. Only JOnTADS, OnTAD, and rGMAP completed the run within a reasonable time frame, while DeTOKI was unable to finish the run within 48 hours on a 2.8 GHz Intel Xeon Processor with 256 GB RAM, and 3DNetMod produced errors despite several attempts with different parameter settings (details of running time in Supplementary Table 1). The number of boundaries identified by JOnTADS, OnTAD and rGMAP were summarized in Supplementary Table 6.

JOnTADS produced a TAD hierarchy that agreed well with visual assessment and fitted better with the patterns in the contact matrix than TADs called by OnTAD and rGMAP (Figure 6a). Moreover, the boundaries identified by JOnTADS and OnTAD had a higher signal of CTCF and Rad21 than those identified by rGMAP, confirming the biological relevance of their identified boundaries (Figure 6b). Additionally, JOnTADS also had the highest average TAD adj-*R*^2^, indicating that its inferred TAD hierarchy explained the variation of the Micro-C contact map much better than rGMAP (Figure 6c).

**Figure 6:**
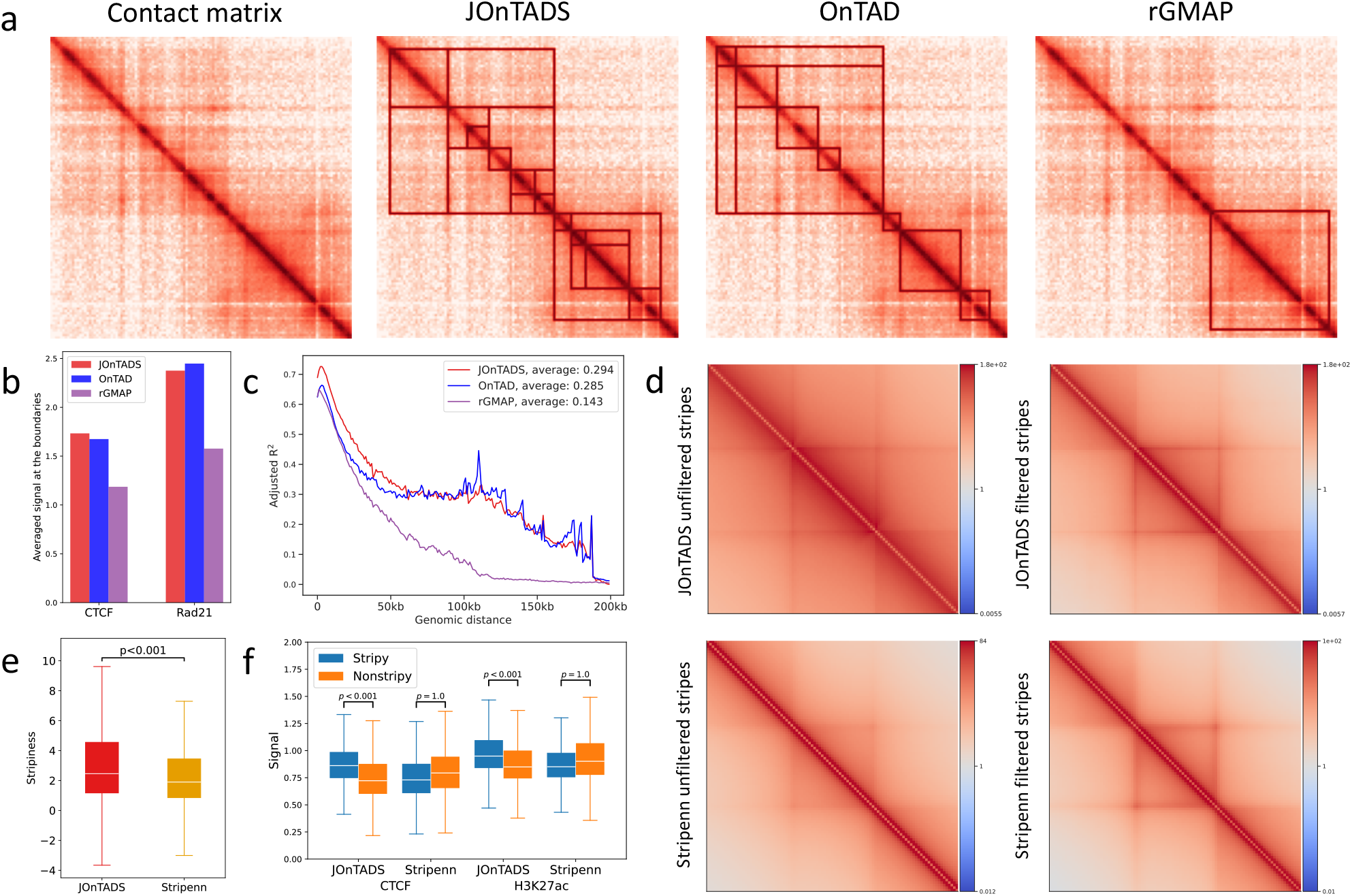
TAD and stripe identification in H1-hESC Micro-C data. **a** TADs identified by JOnTADS, OnTAD and rGMAP, displayed in the view window of the contact matrix at 19.65mb-19.78mb on chromosome 22. **b** Averaged CTCF and Rad21 signals at identified TAD boundaries spanning a 0-200kb region. **c** TAD-adjusted *R*^2^ for TADs identified by JOnTADS, OnTAD and rGMAP. **d** Average epigenetic state signals at identified TAD boundaries. **e** Pileup plots of stripes identified by JOnTADS and Stripenn. Filtered: stripes passing the filter of Stripenn p-values ≤0.1. Unfiltered: all identified stripes. **f** Stripiness for filtered stripes identified by JOnTADS and Stripenn. **g** Distributions of average CTCF and H3K27ac signals within stripy and non-stripy TADs based on filtered stripes identified by JOnTADS and Stripenn. The p-values were calculated using one-sided t-test.

#### JOnTADS effectively identifies stripes and uncovers epigenomic patterns on Micro-C data

We assessed JOnTADS’s performance on stripe identification in this dataset and compared it to Stripenn [43]. Since Stripenn does not support resolutions finer than 5Kb, we performed the comparison on 5Kb data, and JOnTADS was also run on 1Kb data. JOnTADS identified 10,304 stripes, while Stripenn identified 40,734 on the 5Kb data (Supplementary Table 7), with JOnTADS completing the task in just 4.2% of Stripenn’s time (Supplementary Table 1).

Stripes identified by JOnTADS displayed higher enrichment of contact signals against their surrounding genomic regions than those identified by Stripenn in the pileup plots (Figure 6d). After applying Stripenn’s filter criterion to the identified stripes (Stripenn’s p-value *<* 0.1), the filtered stripes from both methods showed similar visual contrast. However, JOnTADS’s filtered stripes had a significantly higher stripiness score (Figure 6e, p-value *<* 0.001 from one-sided t-test), implying higher continuity than Stripenn’s [43].

To investigate the biological relevance of the identified stripes, we analyzed the average CTCF and H3K27ac signals within stripy and non-stripy TADs. Stripy TADs defined by JOnTADs stripes exhibited significantly higher CTCF and H3K27ac signals than non-stripy TADs in both 1Kb (Supplementary Fig. 3) and 5Kb data (Figure 6f), consistent with previous findings on Hi-C and HiChIP data that TADs with stripes tend to have higher accessibility and more active enhancers [43]. In contrast, the stripy TADs defined by Stripenn showed lower average CTCF and H3K27ac signals than non-stripy TADs on 5Kb data (Figure 6f), further validating JOnTADS’s suitability in stripe identifications on Micro-C data.

## Discussion

In recent years, the field of 3D genomics has seen a rapid increase in the amount and the diversity of genome-wide chromatin conformation data. This includes both population and single-cell data, data with fine resolutions, and data measured under various conditions or cell types. These developments have provided opportunities to investigate the molecular processes involved in the formation of fine-scale interaction patterns, examine cell-specific architectural structures, and compare TADs and stripes across different cell types or conditions, thereby deepening our understanding of the role of these architectural features in gene regulation. Despite this progress, there is still a computational barrier to identifying these features robustly and accurately.

We present JOnTADS, a novel method for detecting both TADs and stripes from various types of genome-wide chromatin conformation data. JOnTADS is highly versatile, capable of analyzing both population and single-cell Hi-C data, both low-resolution and high-resolution data, for both single-sample and multiple-sample scenarios. This versatility is achieved through a series of data-driven modeling strategies that accommodate varying levels of data sparsity. With minimal user input required, JOnTADS automatically adapts to different contexts, minimizing the need for user intervention in parameter tuning. As a unified framework, JOnTADS enables the simultaneous study of both architectural features, promoting consistency in analyses. Our comparative analyses demonstrate that JOnTADS consistently produces biologically more relevant identifications across a range of data types than previously developed TAD and stripe callers.

One unique feature of JOnTADS is its capability to jointly construct TAD hierarchies from multiple contact maps. By leveraging combined information and common structures across maps, it effectively reduces unwanted boundary shifts across samples in TAD identification, while highlighting biologically meaningful differences. Our multi-sample analyses on population Hi-C data demonstrate that the TAD hierarchies from JOnTADS’s joint construction align with the expected lineage relationships in hESC differentiation time and follow biologically anticipated TAD formation dynamics during cell phases in mouse G1E-ER4 cells. These results showcase JOnTADS’s effectiveness in comparing samples across different conditions or states to elucidate the roles of chromatin structures in gene regulation.

JOnTADS’s joint analyses also enable robust identification of TLDs in single-cell Hi-C data, effectively addressing challenges posed by data sparsity. This approach reveals insights into the relationship between the frequency that a bin is identified as a TLD boundary across single cells and its biological characteristics in GM12878 Dip-C data [24]. It also yields TLD identifications that more effectively distinguish cell types in human brain prefrontal cortex Hi-C data [54] compared to existing methods. These findings contribute to a deeper understanding of genome organization and regulation at the single-cell level.

Finally, JOnTADS demonstrates high computational efficiency, particularly when handling high resolution data such as Micro-C. These attributes collectively make JOnTADs a highly suitable tool for the comparative analyses of architectural features across a wide array of samples collected by the 3D genomics community, facilitating a better understanding of the mechanisms governing TAD and stripe formation in various cell types and conditions.

## Methods

### Normalization

Let *X*^*raw*^ be the raw Hi-C contact matrix of size *N*, where each bin 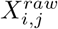 is the interaction frequency between bins *i* and *j*. Let *d* be the genomic distance between the two interacting bins, i.e. *d* = |*i* − *j*|.

To make the interaction frequency comparable across genomic distances and contact matrices, the raw contact frequency in each genomic distance is rescaled to the range of [0, 1] using the min-max normalization,

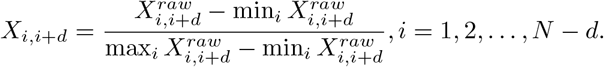

To improve stability, we approximate max_*i*_ 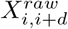 using 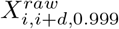, the 99.9^*th*^ percentile of 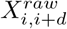. The final normalization equation is

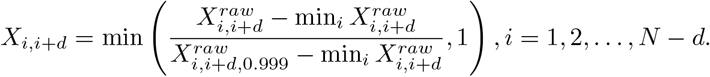

*X*_*i,i*+*d*_ is used as the input for all the subsequent analyses. The notations 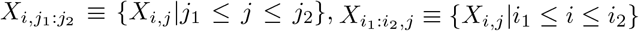, and 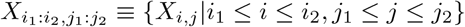, are used to denote a horizontal line at the *i*^*th*^ row spanning *j*_1_ to *j*_2_ columns, a vertical line at the *i*^*th*^ column spanning *j*_1_ to *j*_2_ row, and a rectangle region on a contact matrix, respectively.

### Identification of candidate boundaries

To identify candidate boundaries on a contact matrix, JOnTADS scans the matrix along the diagonal in both directions, using a pair of lines that are parallel to the matrix sides and extend from the matrix diagonal (Figure 1a). At each scanned location, JOnTADS computes the average values of several strongest bins on the line and identifies locations showing significantly different averages from their adjacent neighbors as potential boundaries. Unlike existing insulation-based TAD-calling methods, such as OnTAD [33], which use square-shaped scanning windows to detect regions with substantial contrasts in average signals between the window and its surroundings, JOnTADS’s line-shaped scanning detects lines with large contrasts between the top signals on the line and those on adjacent lines. This method is more sensitive to TADs with pronounced boundary signals but weaker internal signals, which are quite common in sparse data, such as single-cell Hi-C experiments. In contrast, square-shaped windows in such contexts may not effectively detect TADs, as low within-TAD interaction frequencies could reduce overall signal averages. The detailed procedure is described as follows.

To proceed, JOnTADS computes the signals on a line 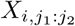 by taking the average contact frequency of its top *K* bins, denoted as 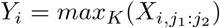, where *max*_*K*_ represents the average of the highest *K* values. Denote the change of signal between two adjacent lines as *d*_*i*_ = *Y*_*i*_ − *Y*_*i−*1_ and the corresponding random variable as *D*. For lines that pass boundaries, *D* tends to have large signal changes with high variability; otherwise, *D* tends to have small changes with low variability. Thus, JOnTADS models *D* using a Gaussian mixture model, 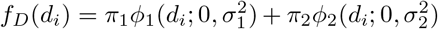, where *π*_1_ + *π*_2_ = 1, *σ*_1_ *> σ*_2_, *ϕ*_1_ and *ϕ*_2_ are normal probability density functions, representing the distributions of signals on lines passing through boundaries and those that do not, respectively. It then estimates *π*_1_, *π*_2_, *σ*_1_ and *σ*_2_ by maximizing the data likelihood, *∏*_*i*_ *f*_*D*_(*d*_*i*_), and identifies the lines with *d*_*i*_ *>* 2*σ*_1_ + *π*_2_ × *σ*_2_ as potential TAD boundaries. These lines show large signal changes between adjacent locations, potentially crossing boundaries. The ends of these lines on the diagonal of the matrix are considered as candidate TAD boundaries.

To identify TADs of various sizes, JOnTADS scans the line pair across contact matrix multiple times, each with a different length. For our analyses, five different lengths are used, evenly spaced within the range of (*T*_*min*_ − 2, *T*_*max*_ − 2), where *T*_*min*_ and *T*_*max*_ are the minimum and maximum TAD sizes in terms of bins specified by users. *T*_*min*_ = 7 and *T*_*max*_ = 200 are used for all of our analyses. Because the regions near the diagonal offer little information for locating boundary positions, the start of all lines is set 2 bins away from the diagonal, and *Y*_*i*_ is computed using *K* = *T*_*min*_ − 2. Then the candidate boundary identified from all line scans are combined into a candidate boundary set, 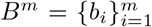, where *b*_*i*_ represents boundary positions indexed according to the order of their genomic locations on the chromosome, i.e. *b*_1_ *< b*_2_ *<* … *< b*_*m*_. To ensure that final boundaries are at least 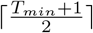 bins apart, the candidate boundaries are decluttered by keeping only the boundary with the highest signal in each neighborhood of 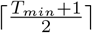 bins (Algorithm 1).

#### Algorithm 1

Prune candidate boundaries

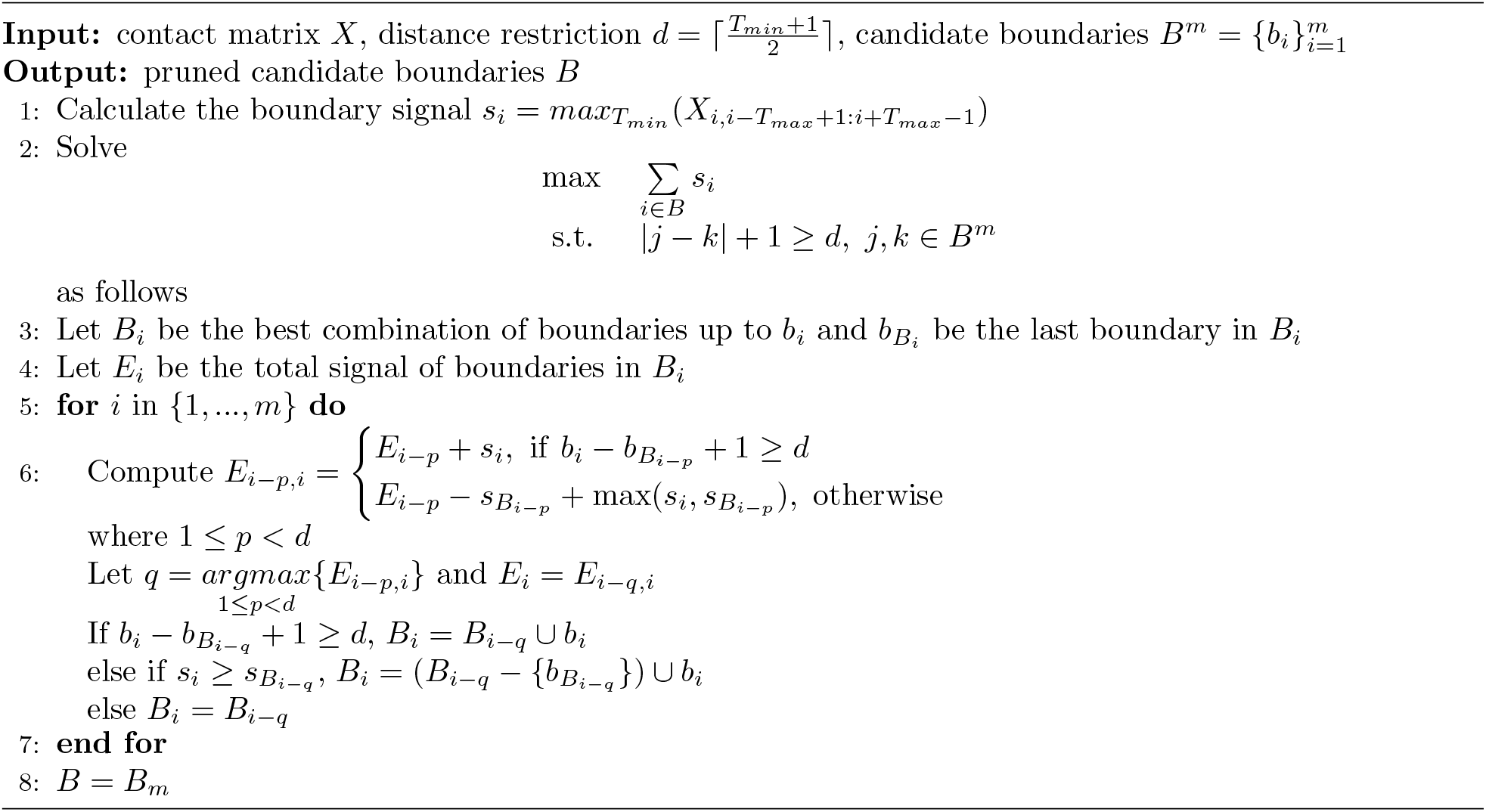

The entire procedure is summarized in Algorithm 2, which outputs the candidate boundary set, 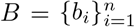.

#### Algorithm 2

Identification of candidate boundaries

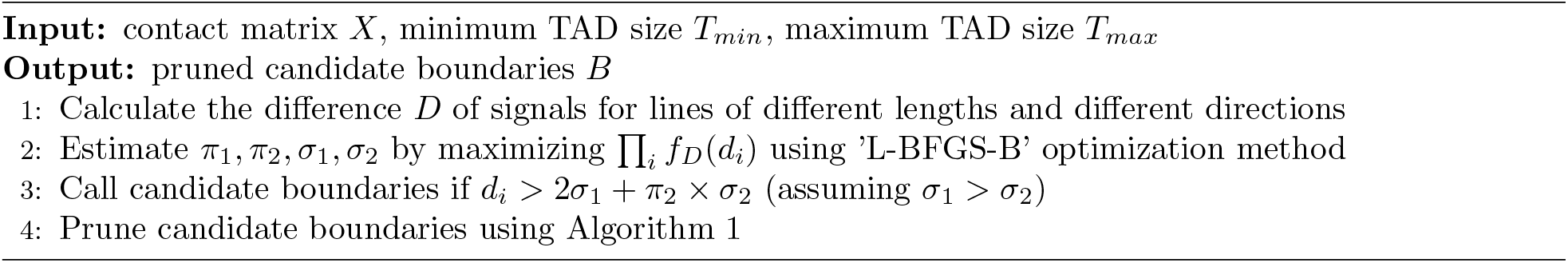

### Construction of TADs from candidate boundaries

#### TAD score

To construct the TAD hierarchy, JOnTADS first assigns a TAD score △ to each candidate TAD region, and then uses a dynamic programming algorithm to determine the optimal hierarchy with the highest overall score. The TAD score is defined as the difference between the mean signal of a diagonal block matrix, excluding any sub-TADs nested inside, and the mean signal of surrounding regions. Specifically,

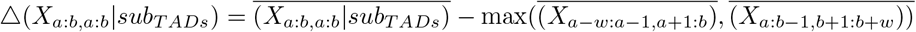

where *w* is the width of the surrounding rectangle region and is defined as

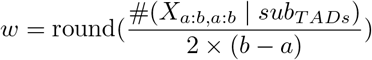

Here, 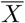 represents the mean signal of a region *X, X*_*a*:*b,a*:*b*_|*sub*_*TADs*_ represents the signals in the region of *X*_*a*:*b,a*:*b*_ excluding the bins within the subTADs, and # means the number of bins within a region. Since the interaction within a TAD is typically higher than that outside of a TAD, a potential TAD is expected to have a positive TAD score.

While both OnTAD and JOnTADS score candidate TAD regions by comparing the mean signal between the candidate TAD and its surrounding region, JOnTADS’s score function offers two advantages over OnTAD’s. First, the function used by JOnTADS contains almost the same number of bins in the surrounding region as in the candidate TAD region, while OnTAD’s function includes about twice as many bins, which can introduce bias due to differences in genomic distance. Second, JOnTADS only considers the mean signal intensity of the region outside of the subTADs, excluding all subTADs, when determining whether a candidate TAD should be called. In contrast, OnTAD considers all subTADs within the candidate TAD. As a result, OnTAD tends to over-call putative regions containing high-intensity inner TADs, while JOnTADS avoids the influence of inner TADs on the calls of outer TADs.

### Dynamic programming procedure for TAD construction

JOnTADS uses a dynamic programming algorithm to construct a TAD hierarchy based on TAD scores. The objective of this procedure is to find a TAD hierarchy that maximizes the sum of TAD scores for the entire hierarchy, i.e.,

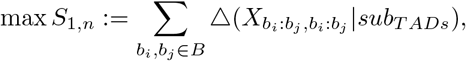

where *S*_1,*n*_ represents the sum of TAD scores for the entire TAD hierarchy between boundaries *b*_1_ and *b*_*n*_ 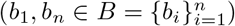.

To determine the TAD hierarchy, JOnTADS calculates TAD scores for the innermost diagonal matrices first and then computes TAD scores for outer diagonal matrices. In light of the observation that the apex region of a TAD often exhibits relatively high signals due to interactions between the flanking bins of the two boundaries, JOnTADS requires that the mean signal for the square region (1/10 of the TAD size) near the apex of a potential TAD must be at least 30% of the mean signal of the entire TAD. Only regions meeting this condition are designated as TADs during the dynamical programming procedure.

To enhance computational efficiency, JOnTADS employs a two-stage dynamic programming procedure, utilizing the fact that the TAD size is constrained by *T*_*max*_. In Stage 1, the algorithm traverses the boundary set and constructs a TAD hierarchy at each boundary, where each boundary serves as the left boundary of a TAD with a size limit of *T*_*max*_. In Stage 2, the algorithm compares different TAD configurations obtained in Stage 1 across the entire contact map, and then selects the optimal configuration using Algorithm 3. Notably, while it is possible to perform the task in Stage 2 by traversing the entire contact map using the same algorithm as in Stage 1, this approach would require a large amount of unnecessary computation for structures larger than *T*_*max*_, resulting in a computational complexity of *O*(*n*^3^), where *n* is the number of candidate boundaries. In contrast, the two-stage approach sidesteps this unnecessary computation, reducing the computational time to O((*T*_*max*_*/T*_*min*_)^2^ × *n*).

#### Algorithm 3

Construction of TADs from candidate boundaries

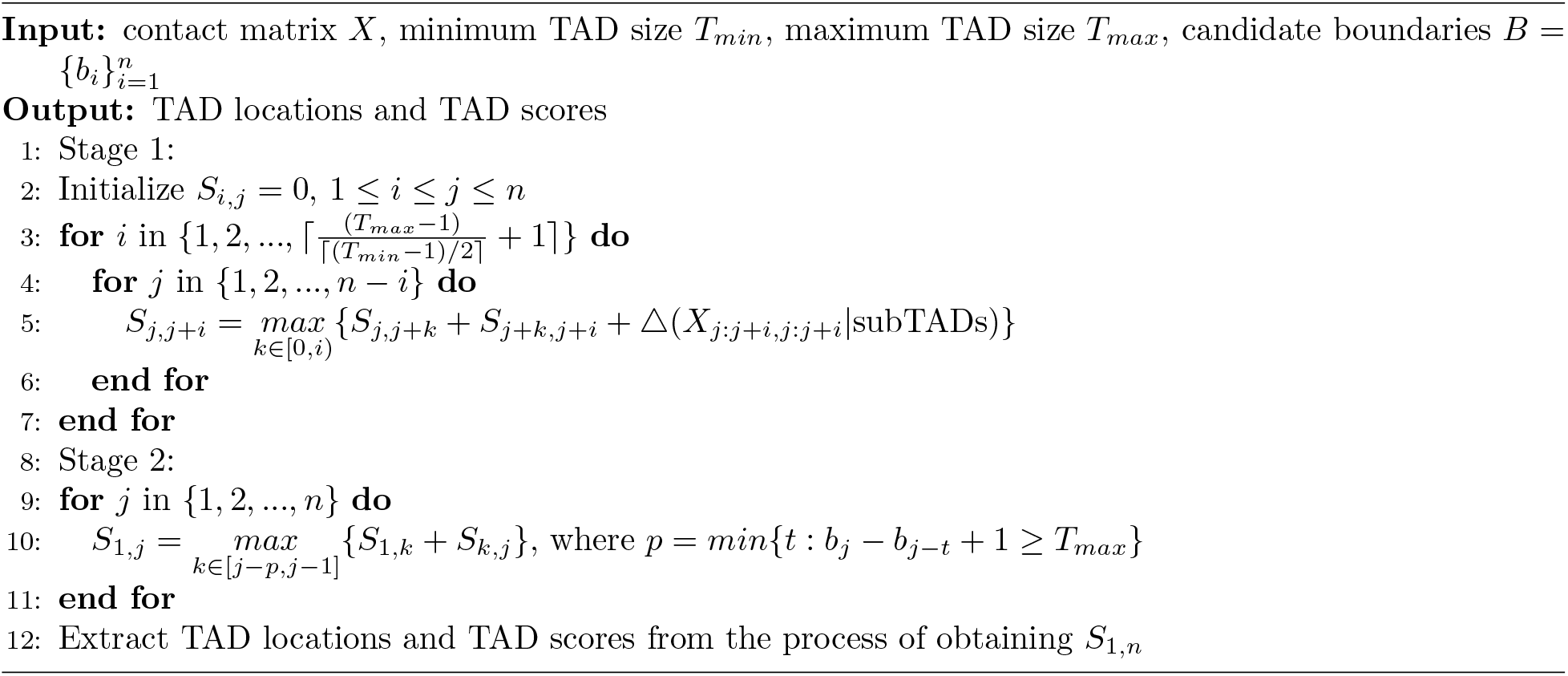

### FDR control for TAD calling

Inevitably, some TADs identified by the dynamic programming procedure may be false positives. To identify these false calls, JOnTADS generates a null matrix devoid of real TADs by shuffling the bins of the contact matrix at each genomic distance and then applies the TAD-calling procedure to the shuffled matrix. The resulting TADs are expected to represent false positives. The TAD scores obtained from the shuffled matrix are used to estimate a TAD-calling threshold that controls the false discovery rate.

However, relying on a single TAD-calling threshold is insufficient, as TAD scores display a non-linear decreasing trend with increasing TAD size (Supplementary Fig. 2a-b). To address this, JOnTADS employs a nonparametric quantile regression model with a non-increasing constraint [55] to estimate the trend. This approach allows us to robustly estimate how TAD size influences TAD scores across different quantiles, effectively capturing the non-increasing trend while mitigating the effects of outliers and heteroscedasticity. To enhance computational efficiency, the model is fitted on a subsample of TADs rather than the entire set. To ensure large TADs are reasonably represented in the subsample, JOnTADS samples 100 TADs from each of three size ranges (0-60^*th*^, 60-90^*th*^, and 90-100^*th*^ percentiles). For each TAD size, the TAD with the largest score is included in the subsample to ensure full coverage of the score range.

The quantile regression model is applied to both the TADs constructed from the original and shuffled matrices. For a given TAD score quantile, the model produces a fitted curve for each matrix. TADs with scores above these curve are considered valid. The False Discovery Rate (FDR) is then computed as

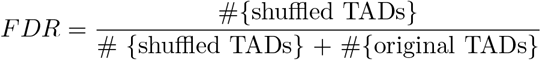

where shuffled TADs and original TADs refer to TADs called in the shuffled and original matrices, respectively.

The TAD score threshold is determined as the lowest quantile that achieves *FDR < α*, where *α* = 0.1 in all of our analyses, using a bisection algorithm. This algorithm iteratively divides the quantile interval in half, starting with an initialization at the 0.5 quantile, fits the model with the quantile at the midpoint, computes the FDR, and then selects the subinterval where the sign of *FDR − α* changes until reaching a precision of 0.01 quantile. Only boundaries of TADs that surpass this threshold are retained. The final TADs are then reconstructed from these retained boundaries by rerunning the dynamic algorithm (Algorithm 3). Algorithm 4 summarizes the entire procedure for a single contact matrix input.

#### Algorithm 4

JOnTADS procedure for a single contact matrix

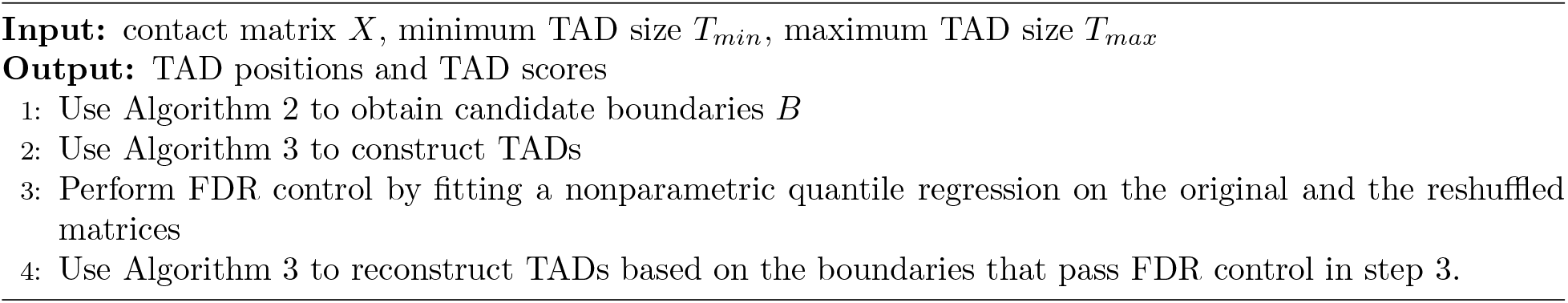

### Joint TAD calling algorithm

To jointly call TADs from multiple contact matrices, JOnTADS first identifies boundaries for each contact matrix using Algorithm 2 and then takes the union of these boundaries across all matrices. For each boundary in the union, JOnTADS computes the average of boundary signals across all matrices and declutters the boundaries based on these average signals using Algorithm 1. For each individual matrix, a subset of decluttered boundaries with the strongest signals in the original matrix is retained. Specifically, denote the original boundary set on the matrix *i* as *B*_*i*_, the decluttered boundary union set as *B*^*d*^, and their corresponding cardinalities as |*B*_*i*_| and |*B*^*d*^|, respectively. Let |*B*_*max*_| = *max*_*i*_(|*B*_*i*_|) be the number of boundaries in the contact matrix with the largest number of boundaries among all matrices. Then the matrix *i* keeps 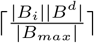 number of boundaries in *B*^*d*^ that have the highest boundary signals on the matrix *i*, where ⌈· ⌉ is the ceiling function. The resulting boundaries are used to construct TAD hierarchies for each matrix using Algorithm 4, which consolidates boundaries that have slightly shifted locations across matrices. Figure 1b provides an overview of the joint TAD calling procedure.

### Stripe identification

To identify stripes on a contact matrix, JOnTADS first normalizes the matrix as outlined in Methods-Normalization, and subsequently scans the diagonal of the normalized matrix to detect the presence of stripes at each location and determine their lengths. We define a stripe as a line segment of 1-bin wide that starts at the diagonal of the contact matrix and extends either horizontally or vertically. Here we describe the algorithm for identifying horizontal stripes; the algorithm for vertical stripes follows similarly.

Denote *X*_*i,i*:*i*+*k*_ as a horizontal stripe of length *k* that starts at *X*_*i,i*_ and ends at *X*_*i,i*+*k*_. We set the maximum and minimum lengths of a stripe to be the same as those of a TAD, namely, *T*_*max*_ and *T*_*min*_, respectively. To detect stripes, JOnTADS searches for line segments that have enriched signals compared to their adjacent neighbors at the same genomic distance. Define a hit set *H*_*i*_ to record the bins on a line originating from diagonal location (*i, i*) that meet this criterion,

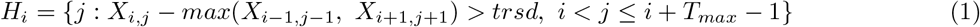

where *trsd* is a threshold parameter representing the minimum sufficient contrast between *X*_*i,j*_ and its adjacent neighbors at the same genomic distance to qualify *j* as a hit. By default, *trsd* is set as 0.1. Only the line that has at least 1*/*10 of its bins in the hit set, i.e., 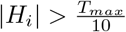, is considered a putative stripe.

Next, JOnTADS assigns scores to each potential stripe and determines its off-diagonal termination point. Genuine stripes typically exhibit a prevalence of ‘hit’ bins (i.e., those with high signals) near the diagonal. As the distance from the diagonal increases, the frequency of these ‘hit’ bins gradually diminishes. A higher proportion of ‘hit’ bins within a potential stripe signifies a greater likelihood of it being a genuine stripe. The shift from predominantly “hit” bins to predominantly ‘miss’ bins signifies the conclusion of a stripe.

To quantify the confidence of a stripe, we develop an enrichment score inspired by the gene set enrichment statistic [44], which assesses the significance of a gene set within a ranked gene list. Let *hit*_*i,j*_ denote the strength of a hit at position (*i, j*), defined as *hit*_*i,j*_ = 2*X*_*i,j*_ *−* (*X*_*i−*1,*j−*1_ + *X*_*i*+1,*j*+1_). Then we compute the cumulative fraction of ‘hit’ bins, weighted by their signal strength, and the cumulative fraction of ‘miss’ bins up to a given position *l* from the diagonal,

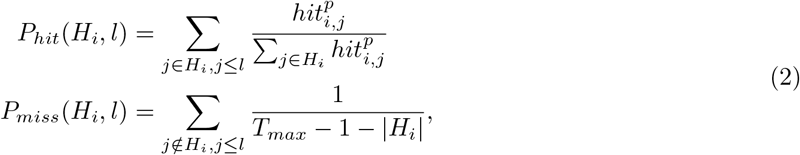

where *p* is typically set as 1. In the context of stripe identification, ‘hit’ bins tend to dominate “miss” bins near the diagonal and gradually decrease in prevalence when being away from the diagonal. Thus, *P*_*hit*_(*H*_*i*_, *l*) − *P*_*miss*_(*H*_*i*_, *l*) initially increases with the increase of *l* and subsequently decreases after ‘miss’ bins become more prominent than ‘hit’ bins. To determine the termination point of a stripe, we seek the position that maximizes the difference in cumulative fractions of ‘hit’ and ‘miss’, denoted as *P*_*hit*_(*H*_*i*_, *l*) − *P*_*miss*_(*H*_*i*_, *l*). The termination point is represented as *l*_*i*_ = *argmax*_*l*_(*P*_*hit*_(*H*_*i*_, *l*) − *P*_*miss*_(*H*_*i*_, *l*)). Additionally, we introduce the stripe’s enrichment score (ES) as *ES*_*i*_ = *max*_*l*_(*P*_*hit*_(*H*_*i*_, *l*) − *P*_*miss*_(*H*_*i*_, *l*)), akin to the gene set enrichment score. For a candidate, a high value of *ES*_*i*_ suggests a stronger signal compared to neighboring regions, while a value near zero implies similarity to the background signal. Similar to the gene set enrichment score, *ES*_*i*_ reduces to the standard Kolmogorov–Smirnov statistic when *p* = 0. To ensure the enrichment of “hit” bins at proximal distances, we require that the cumulative average deviation of a stripe between *P*_*hit*_(*H*_*i*_, *l*) and *P* (*H, l*) exceeds a positive threshold, i.e. 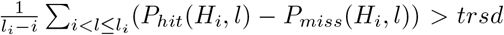. Here we used the same threshold as the one used in equation 1. Algorithm 5 outlines the procedure for identifying horizontal stripes in detail.

#### Algorithm 5

JOnTADS stripe identification

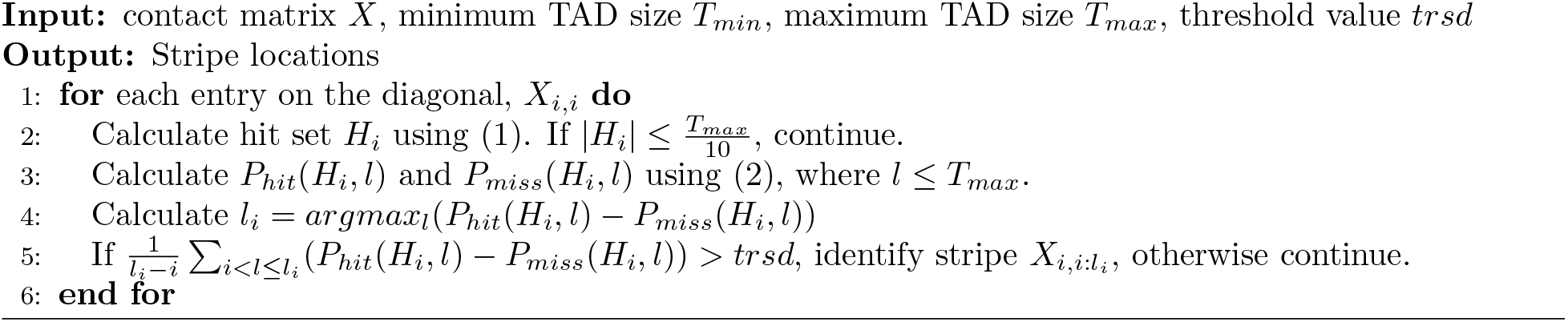

### Metrics for characterizing similarity and distinctiveness of TAD or TLD structures between a pair of cells

#### Jaccard Index for TADs

We used Jaccard Index (JI) to quantify the similarity between two sets of TAD structures. Given the significant differences in variation between population and single-cell data, we define two versions of Jaccard Index for TADs: one for population Hi-C and micro-C data and another for single cell Hi-C data, referred to as JIT and scJIT, respectively.

(1) Jaccard Index for TADs for population data (JIT)
Given two sets of TADs, *T*_1_ and *T*_2_, the Jaccard Index for TADs for population data is defined as

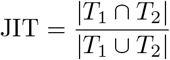

where *T*_1_ ∩ *T*_2_ contains TADs with exactly the same left and right boundaries. That is, suppose *X*_*a*:*b,a*:*b*_ ∈ *T*_1_, *X*_*c*:*d,c*:*d*_ ∈ *T*_2_, if *a* = *c* and *b* = *d*, then *X*_*a*:*b,a*:*b*_ and *X*_*c*:*d,c*:*d*_ belong to *T*_1_ ∩ *T*_2_. A higher Jaccard Index implies a higher degree of similarity between *T*_1_ and *T*_2_.

(2) Jaccard Index for TADs for single cell data (scJIT)

Single-cell Hi-C data is highly variable due to the heterogeneous chromatin conformations across cells. Consequently, the standard Jaccard Index often yields near-zero values when measuring TAD overlap between single-cell Hi-C matrices, making it difficult to distinguish the performance of different methods. To address this, we developed a modified Jaccard Index for comparing TLS overlaps across samples, denoted as scJIT. Unlike the standard JI, which only considers fully overlapped TADs as intersections and ignores partial overlaps, the modified JI includes both fully and partially overlapping TADs in the intersecting set. It accounts for the overlap of both regions and boundaries, weighing the degree of overlap. Specifically, let *T*_1_ and *T*_2_ represent two sets of TADs, then scJIT is defined as

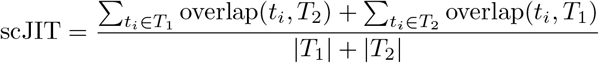

where | · | represents the cardinality of the corresponding set and

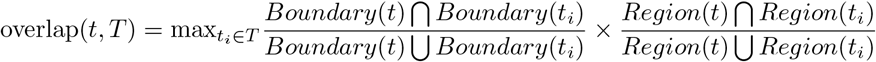

Here *Boundary*(*t*) and *Region*(*t*) are the sets of bins in the boundaries and the TAD region of TAD *t*, respectively. The first term in overlap(*t, T*) represents the Jaccard Index of boundaries, and the second term represents the Jaccard Index of TAD regions. This formula can be viewed as a special case of the soft Jaccard index in [56]. Given the same input, scJIT is always greater than JIT.

### Common TLS Enrichment Score (CTES) and Differential TLS Enrichment Score (DTES)

To characterize TLSs that are commonly or specifically identified in a pair of single cells, we devised two metrics: the Common TLS Enrichment Score (CTES) and the Differential TLS Enrichment Score (DTES). CTES measures the signal strength for TLSs with identical boundaries in both samples, while DTES quantifies the signal contrast for the remaining sample-specific TADs. A robust caller should achieve high score in both metrics, reflecting strong signal enrichment for common TLSs and strong signal contrast across samples for sample-specific TLSs.

To compute these metrics, we first normalize the contact frequency within each TLS by dividing the total contact frequency within the domain by the average total contact frequency of a same sized square diagonal region on the same chromosome, excluding the diagonal itself. The normalized value represents signal enrichment that is comparable across samples and TADs of different sizes, addressing the high variability in single-cell data. CTES is defined as the average enrichment across samples for common TADs. For each sample-specific TLS, the enrichment difference between two samples is calculated for the region covered by the TLS, and DTES is defined as the average of these differences across all sample-specific TLSs.

Specifically, let *T*_*c*_ = (*t*_1_, …, *t*_*k*_) be the set of TADs that are commonly identified across samples A and B, and 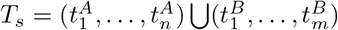 be the set of sample specific TADs. Let *E*_*j*_(*t*_*i*_) be the enrichment of TAD *t*_*i*_ on sample *j* and *R*(*t*_*i*_) be the region with the same coordinates as *t*_*i*_, then

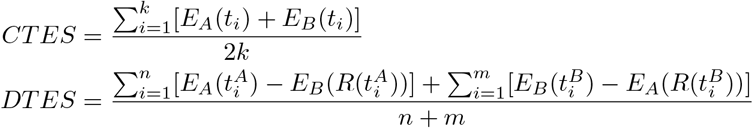

### Comparison of TAD calling and stripe calling methods

We analyzed 40kb bulk GM12878 data [28], 40kb bulk hESC lineage data [46], 10kb bulk G1E-ER4 cell cycle data [18], 40kb Dip-C GM12878 data [23], 10kb single-nucleus methyl-3C sequencing (sn-m3C-seq) data [54], and 5kb and 1kb Micro-C H1-hESC data [19].

We compared JOnTADS with three hierarchical TAD calling models (OnTAD, rGMAP, and 3DNetMod) and a single cell TAD calling method (deTOKI). JOnTADS and deTOKI were run on raw contact matrices for all datasets. OnTAD, rGMAP, and 3DNetMod were run on normalized matrices, as required by each method. We generated normalized matrices using the Knight-Ruiz method [57] for Micro-C H1-hESC data and the ICE method [58] for all other datasets. For the bulk G1E-ER4 cell cycle data from [18], we used the published 3DNetMod results (downloaded from https://www.nature.com/articles/s41586-019-1778-y, under the label “most merged version”), rather than running 3DNetMod in-house. For datasets with multiple samples (i.e. bulk hESC lineage data, bulk G1E-ER4 cell cycle data, and Dip-C GM12878 data), we applied JOnTADS to all contact matrices jointly.

We used the recommended parameter settings for all methods during data analyses. Due to the large size of the Micro-C data for bulk H1-hESC (raw contact matrix is 50819 × 50819 in size, taking 4.9GB of memory), we restricted our analysis to chromosome 22. For the remaining datasets, we analyzed all autosomes (i.e. chromosomes 1-22 for bulk GM12878, bulk hESC lineage, and GM12878 Dip-C data; and chromosomes 1-19 for mouse bulk G1E-ER4 cell cycle data). To assess running time, we compared the performance of the methods on a single chromosome. For bulk human ES cell lineage data, we performed hierarchical clustering using ‘average’ linkage based on the Jaccard Index for TADs.

To identify architectural stripes, we used the Stripenn settings from its Github example run (https://github.com/ysora/stripenn). Specifically, we executed the following command: `stripenn compute --cool 5000.cool --out output dir/ -k 4 -m 0.95, 0.96, 0.97, 0.98, 0.99` where 5000.cool represents the 5Kb resolution Micro-C data. To compare our results with Stripenn [43], we applied Stripenn’s score function to our identified stripes. This function provides two values for each stripe: a p-value reflecting the significance of the difference between the stripe and its surrounding regions, and a stripiness score measuring the continuity of signals within the stripe. Similar to the approach in [43], we used a p-value cutoff of 0.1 to filter our stripes. For the Micro-C H1-hESC dataset [19], the Stripenn score function encountered errors when applied to our results for chromosomes 2, 11, 16, and 20, thus only the results for the rest 18 chromosomes were reported. For the GM12878 dataset from [28], Stripenn score function runs into error when applied to our results for chromosome 6, thus only the results for the rest 21 chromosomes were reported.

### IDEAS state enrichment

We used IDEAS [53], an epigenomic annotation system that integrates data from multiple epigenomic experiments to infer chromatin states based on combinatorial patterns of histone modifications, to obtain epigenetic states for the bulk/Dip-C GM12878 dataset. The IDEAS states of GM12878 cell type, obtained in [52], were used for our analysis. In the study by [52], the entire genome was divided into 200bp intervals, and IDEAS was applied to five histone marks of 127 cell types from the Roadmap Epigenomics Consortium [59] to determine the epigenetic state for each interval. A total of 20 distinct epigenomic states were obtained. To compute the signal strength of each IDEAS state for each Hi-C bin, we counted the number of 200bp intervals annotated as that state within the bin. We then assessed the enrichment of each IDEAS state in a Hi-C bin by computing the ratio of its signal strength to the genome average, i.e.

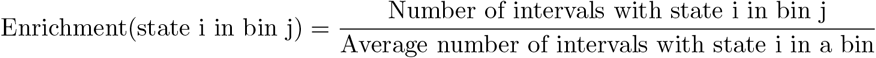

We classified the 20 IDEAS states into active or repressed states based on the absence or presence of H3K27me3 and H3K9me3, both of which play repressed roles in the genic and nongenic regions of metazoan genomes [60]. As shown in Figure 1A in [52], the states, 2_TxWk, 4_Enh, 5_Tx, 6_EnhG, 8_TssAFlnk, 10_TssA, 14_TssWk, and 17_EnhGA, lacked H3K27me3 or H3K9me3 signals, while the other states showed at least one of H3K27me3 and H3K9me3 signals. So we classified the former as active states, and the latter as repressed states.

For the Micro-C H1-hESC dataset, we used the epigenetic state annotations generated by ChromHMM [61] and published in EpiMap [62]. These annotations were based on the same genome build (hg38) as the Micro-C dataset. The ChromHMM analysis resulted in 18 distinct epigenetic states. We calculated the enrichment using the same formula as used for IDEAS.

## Data availability

Contact matrix data: The bulk GM12878 Hi-C data is from [28] (GEO accession number: GSE63525). The bulk hESC lineage Hi-C data is obtained from [46] (GEO accession number: GSE52457). The G1E-ER4 cell cycle data comes from [18] (GEO accession number: GSE129997). The Dip-C GM12878 data is from [23] (GEO accession number: GSE117876). The processed single-nucleus methyl-3C sequencing (sn-m3C-seq) data is obtained from https://github.com/dixonlab/scm3C-seq [54]. The bulk Micro-C H1-hESC data [19] is obtained from the 4D Nucleome Data Portal (4DNFI2TK7L2F.hic).

Epigenomic data: ChIP-seq data of GM12878 and H1-hESC cell lines are downloaded from ENCODE project https://www.encodeproject.org/. For GM12878, CTCF accession is ENCFF312KXX, Rad21 accession is ENCFF550QIX, Smc3 accession is NCFF235BXX, and H3K27ac accession is ENCFF180LKW. For H1-hESC, CTCF accession is ENCFF473IZV, Rad21 accession is ENCFF002NBT, and H3K27ac accession is ENCFF919FBG. All files are downloaded in ‘bigWig’ format and processed by ‘bigWigAverageOverBed’ downloaded from UCSC Genome Browser.

Epigenetic states: The IDEAS epigenetic state of the GM12878 cell type is downloaded from http://bx.psu.edu/~yuzhang/Roadmap_ideas/ [52]. The file name is ‘test1.114.bb’ [53, 52].

## Code availability

JOnTADS package is available at https://github.com/qunhualilab/JOnTADS under MIT license.

## Acknowledgements

Q.L. acknowledges support from NIH grant R01GM109453 and R.C.H acknowledges support from NIH R24DK106766. Q.L. acknowledges support from seed grants of the Institute for Computational and Data Sciences and Consortium on Substance Use and Addiction at the Pennsylvania State University. This study used computational resources provided by the Institute for Computational and Data Sciences at the Pennsylvania State University.

## Author contributions

Q.L. conceived and supervised the study. Q.Z. and Q.L. developed the methods. Q.Z. performed the analyses and visualized the results with input from Q.L., R.C.H., Y.Z. and G.X. Q.L. and Q.Z. wrote the manuscript with input from R.C.H.

