## Supplementary methods, figures, and tables for "JOnTADS: a unified caller for TADs and stripes in Hi-C data"

#### Parameter choice

When using JOnTADS, users only need to specify two parameters, the maximum and minimum sizes of TADs in bins. These choices should be informed by the data resolution and typical TAD sizes. In our analyses, we consistently set the minimum and maximum TAD sizes to 7 and 200 bins, respectively. To identify candidate boundaries, we use 5 lines to scan the diagonal. We control FDR at 10% for TAD calling in all our analyses. While users have the flexibility to adjust these parameter values, our extensive analyses show that the chosen settings are suitable for various data types tested.

#### Time complexity

##### TAD identification

JOnTADS's TAD identification algorithm comprises two parts, boundary identification and TAD construction. In the first part, it scans the contact matrix using a pair of line segments multiple times, each time with different lengths. Since the maximum length of the line is no more than $T_{max}$, each scan has a time complexity of $O(N \times T_{max})$, where $N$ is the width of the contact matrix. The subsequent step of fitting Gaussian mixtures to identify boundaries is computationally negligible. The boundary decluttering algorithm (Algorithm 1 in Methods) operates linearly with respect to the number of candidate boundaries, i.e., $O(n)$, where $n(n << N)$ is the size of the boundary set. Thus the overall time complexity for this part is $O(N \times T_{max})$.

The second part, TAD construction, is done using a two-stage dynamic programming algorithm (Algorithm 3 in the main text). In Stage 1, TAD scores are computed for TADs smaller than $T_{max}$. It starts from the inner most diagonal matrices and then progresses towards the outer layers. When evaluating TAD scores for outer matrices, all inner boundaries are traversed, considering all possible configurations of TAD structures. The time complexity is

$$time\ complexity = \sum_{i=1}^{O(T_{max}/T_{min})} O((n-i) \times i) = O((T_{max}/T_{min})^2 \times n)$$

where $i$ is the number of inner boundaries and $n$ is the size of boundary set. This complexity can be derived by analyzing Algorithm 3: $(n-i)$ comes from the for-loop in row 4, $i$ comes from the max operation in row 5, and $\sum_{i=1}^{O(T_{max}/T_{min})}$ comes from the for-loop in row 3. In Stage 2, JOnTADS finds the optimal TAD hierarchy for the entire contact matrix by traversing the boundary set and iteratively determining the optimal configuration between the first boundary and the $j^{th}$ boundary $(j = 1, \ldots, n)$. Because the boundaries are spaced at least $(T_{min}+1)/2$ bins apart (owing to the decluttering step in Stage 1), for each $j$, at most $O(T_{max}/T_{min})$ possible configurations need to be compared to find the optimal configuration (see calculation of $p$ in Algorithm 3 row 10). Consequently, the time complexity for Stage 1 is $O(T_{max}/T_{min} \times n)$, dominated by Stage 1's complexity. Although there is no guarantee that $(T_{max}/T_{min})^2 \times n > N \times T_{max}$, empirically, we observe that the TAD construction part consistently consumes more time than the boundary identification part.

The FDR control procedure in JOnTADS involves fitting the nonparametric quantile regression using a quadratic programming algorithm. This computational complexity of this step is polynomial with the number of putative TADs. Since our implementation uses a subsample of around 400 TADs to fit the curve, the computational complexity is polynomial with respect to this subsample size, i.e. $O(Poly(400))$. A bisection search is then performed to find the quantile threshold that achieves the desired FDR control. The bisection search is executed 10 times to achieve an accuracy of $1/2^{10} < 0.001$ in the quantile threshold. Therefore, the running time for the bisection search is $O(10 \times Poly(400))$.

Overall, the total time complexity for TAD identification can be expressed as $O(max((T_{max}/T_{min})^2 \times n), 10 \times Poly(400)))$. If the input contact matrix is small, the quadratic programming step primarily determines the running time; otherwise, the TAD construction part becomes the dominant factor.

##### Stripe identification

The process of identifying stripes involves scanning the matrix diagonal and comparing adjacent bins within a distance of $T_{max}$ from the diagonal to obtain the hit set and compute the ES statistics. Therefore, the time complexity of this step is $O(N \times T_{max})$.

### Supplementary Figures

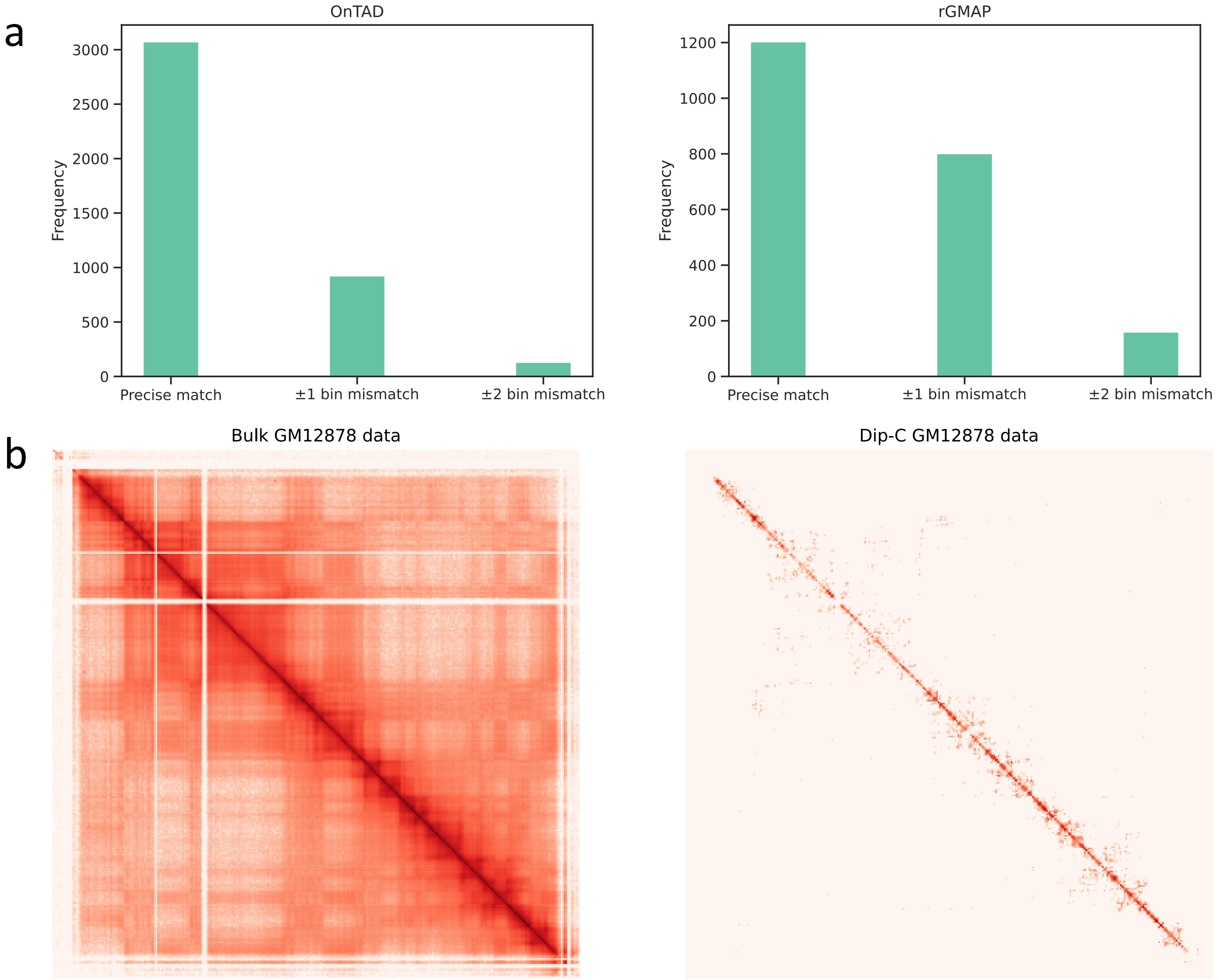


**Supplementary Figure 1**: Boundary matching analysis and Hi-C contact matrices from bulk-cell and Dip-C data. **a** The numbers of matched and shifted boundaries on a pair of human ES cell Hi-C replicate samples. Boundaries are identified by OnTAD (Left) and rGMAP (Right). **b** Heatmaps of the (logarithm transformed) contact matrices for bulk-cell GM12878 (left) and Dip-C GM12878 (right) under 40kb resolution. The region spans from 0 to 13.6million base pairs, i.e. 340 bins, on chromosome 1.

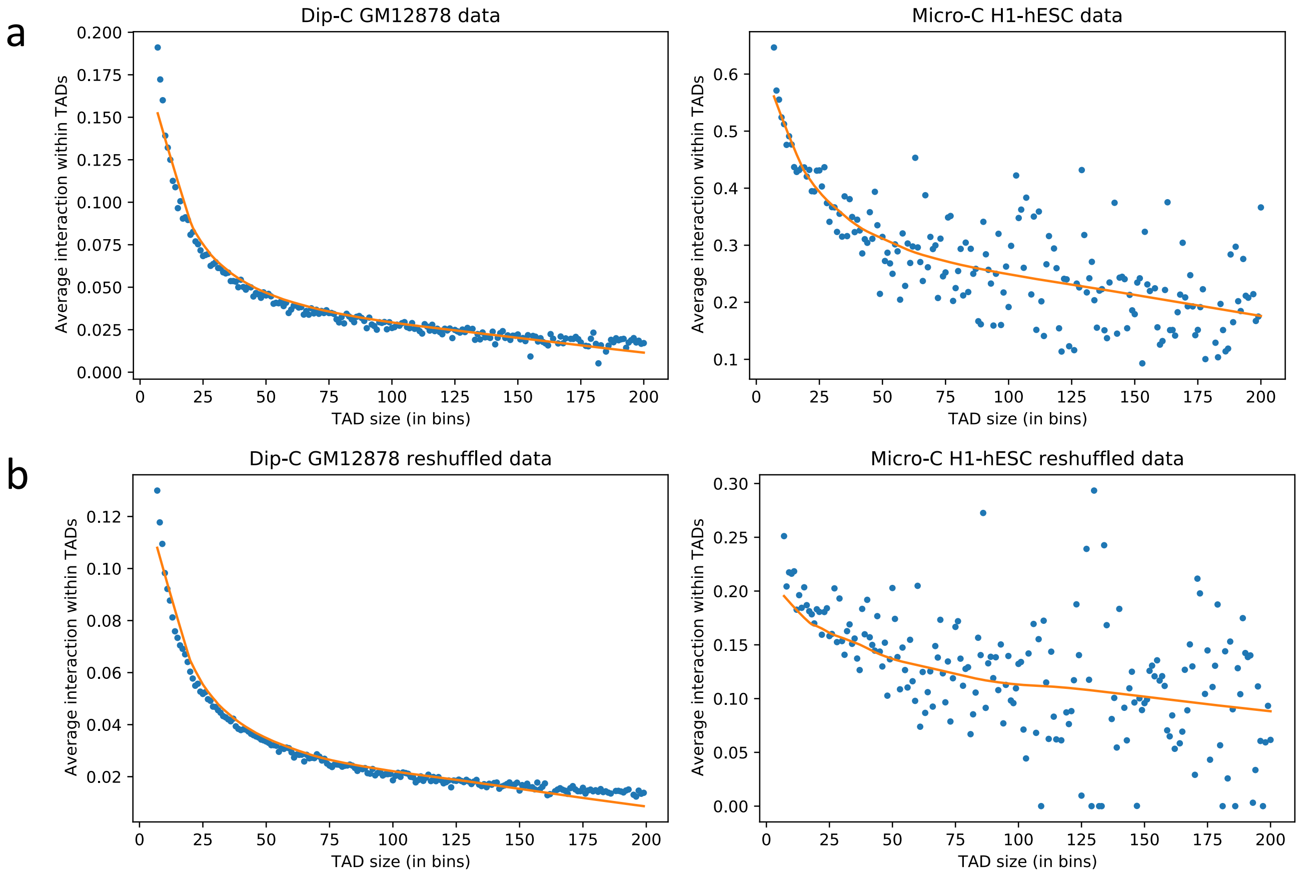


**Supplementary Figure 2**: Relationship between average interaction within TADs and TAD size. **a** Relationship between average interaction within TADs and TAD size for the TADs constructed from the contact matrices before FDR control. **b** Relationship between average interaction within TADs and TAD size for the TAD constructed from reshuffled contact matrices. Loess curves are fitted on the average interaction within TADs at each TAD size (dots). Left: Autosomes of cell02 in the Dip-C GM12878 data; Right: Chromosome 22 in Micro-C H1-hESC data.

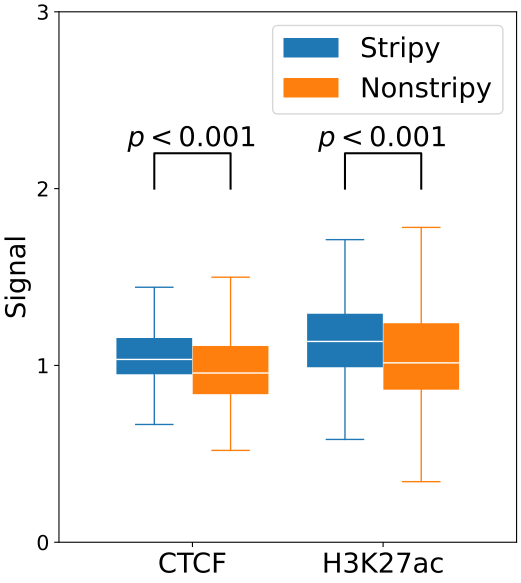


**Supplementary Figure 3**: CTCF and H3K27ac signals at the boundaries of stripy and non-stripy TADs based on stripes identified by JOnTADS on Micro-C data at the 1kb resolution. The stripes are filtered to include only the top 200 stripes with the highest mean signal. The p-values were calculated using one-sided t-test.

### Supplementary Tables

| | Data<br>Programming language | resolution | matrix width | JOnTADS<br>Python | OnTAD<br>C++ | 3DNetMod<br>Python | rGMAP<br>R | deTOKI*<br>Python | deDoc2<br>Java | Higashi<br>Python | scHiCluster<br>Python | Stripenn<br>Python |
|---|---|---|---|---|---|---|---|---|---|---|---|---|
| TAD call | bulk GM12878 chr1 | 40kb | 6232 | 64.5 | 7.4 | 14786.7 | 40.0 | 151.5 | | 3.2* | | |
| | Dip-C GM12878 chr1 (14 cells total) | 40kb | 6232 | 837.3 | 128.4 | | 560.31 | 1862.1 | 300 | > 172800** | 827.2 | |
| | Lee's data chr1 (70 cells total) | 40kb | 6232 | 4049.4 | 665.6 | | 2422.3 | 8202.2 | 1588 | > 172800** | 4427 | |
| | G1E-ER4 cell cycle chr1 (5 stages total) | 10kb | 19720 | 739.3 | 310.7 | 60151.8 | 846.6 | 6899.3 | | | | |
| | Micro-C H1-hESC chr22 | 1kb | 50819 | 322.1 | 367.5 | NA | 550.6 | > 172800 | | | | |
| Stripe call | bulk GM12878 chr1 | 5kb | 49851 | 62.5 | | | | | | | | 3216.8 |
| | Micro-C H1-hESC chr1 | 5kb | 49792 | 73.8 | | | | | | | | 1767.7 |
| | Micro-C H1-hESC chr22 | 1kb | 50819 | 83.0 | | | | | | | | NA |

**Supplementary Table 1**: Running time (seconds) on 2.8 GHz Intel Xeon Processor with 256 GB RAM. *: Only boundaries, no TAD. **: including imputation step, which was run on Google Colab using A100.

| | JOnTADS | OnTAD | 3DNetMod | rGMAP | deTOKI | deDoc2 | Higashi | scHiCluster |
|---|---|---|---|---|---|---|---|---|
| GM12878 bulk cell Hi-C data | | | | | | | | |
| TADs | 4640 | 5993 | 15069 | 2485 | NA | 4893 | NA | NA |
| TAD boundaries | 3576 | 4269 | 15717 | 2285 | 2996 | 7928 | 3579 | NA |
| GM12878 Dip-C data | | | | | | | | |
| TLD | 1637 | 4482 | NA | 2980 | NA | 8152 | 931* | 3710 |
| TLD boundaries | 1844 | 4814 | NA | 2726 | 3061 | 13839 | 3426 | 7419 |

**Supplementary Table 2**: The number of identified boundaries, TADs, and TLDs in the GM12878 bulk cell Hi-C and Dip-C data. The averages across 14 single cells are reported for Dip-C data. The number of bins on the autosomes in the genome is 72,036. *: number of TLDs identified by Higashi+TopDom.

| | prometa | ana-telo | early-G1 | mid-G1 | late-G1 |
|---|---|---|---|---|---|
| JOnTADS | **0.988** (2619) | **0.989** (2320) | **0.991** (2040) | **0.992** (1097) | **0.993** (1759) |
| OnTAD | 0.991 (298) | 0.992 (843) | 0.993 (1332) | 0.996 (1451) | 0.997 (5583) |
| 3DNetMod | 0.995 (272) | 0.995 (901) | 0.997 (1711) | 0.998 (2226) | 0.999 (7810) |
| rGMAP | 0.991 (46) | 0.995 (129) | 0.996 (398) | 0.997 (808) | 0.998 (3025) |
| deTOKI | 0.993 (6) | 0.997 (7) | 1.000 (64) | 0.998 (313) | 0.999 (2250) |

**Supplementary Table 3**: The ratio of insulation scores at boundary positions relative to their adjacent ($\pm 1$) bins for the boundaries identified in the late-G1 phase in the mouse G1E-ER4 cell dataset. Boundaries are stratified by their time of emergence in the cell cycle. Numbers in parentheses denote the count of boundaries in each category. A smaller ratio indicates a larger reduction of insulation score at the boundaries from their adjacent bins, i.e. a stronger insulation.

| | bulk | cell02 | cell03 | cell05 | cell06 | cell07 | cell09 | cell10 |
|---|---|---|---|---|---|---|---|---|
| counts | 3517453218 | 614620 | 722140 | 670374 | 697558 | 797815 | 857446 | 817768 |
| | | cell11 | cell12 | cell13 | cell14 | cell15 | cell16 | cell17 |
| counts | | 1071257 | 984676 | 1064790 | 1018365 | 958032 | 939094 | 952856 |

**Supplementary Table 4**: Read counts of GM12878 bulk Hi-C data and Dip-C data. The cell labels in Dip-C data follow those in Dip-C paper.

| Boundary sharing frequency | JOnTADS | OnTAD | rGMAP | deTOKI | deDoc2 | Higashi+TopDom | scHiCluster |
|---|---|---|---|---|---|---|---|
| 1 | 5276 | 20687 | 17430 | 19259 | 12199 | 1590 | 18534 |
| 2 | 2788 | 9974 | 5847 | 6694 | 12929 | 6561 | 12062 |
| 3 | 1517 | 4220 | 1855 | 2047 | 11174 | 2764 | 6926 |
| 4 | 886 | 1764 | 497 | 626 | 8200 | 1248 | 3674 |
| 5 | 509 | 722 | 152 | 183 | 5562 | 499 | 1924 |
| 6 | 282 | 302 | 35 | 71 | 3559 | 239 | 1115 |
| 7 | 170 | 126 | 12 | 12 | 2173 | 94 | 569 |
| 8 | 95 | 47 | 4 | 7 | 1285 | 62 | 319 |
| 9 | 41 | 13 | 2 | 5 | 756 | 18 | 150 |
| $\geq 10$ | 30 | 21 | 31 | 3 | 740 | 17 | 126 |

**Supplementary Table 5**: Distribution of boundary sharing frequency in GM12878 Dip-C data, where boundary sharing frequency refers to the number of cells in which a boundary is identified.

| | JOnTADS | OnTAD | rGMAP |
|---|---|---|---|
| Boundary number | 1501 | 1704 | 641 |
| TAD number | 1872 | 2254 | 673 |
| CTCF signal | **1.733** | 1.674 | 1.186 |
| Rad21 signal | 2.375 | **2.448** | 1.577 |
| Average TADadj-$R^2$ | **0.294** | 0.285 | 0.143 |

**Supplementary Table 6**: The number of identified boundaries and TADs on Micro-C H1-hESC chromosome 22 data. The average CTCF and Rad21 signal at the boundaries and the average TADadj-$R^2$ over 0-200kb span are also reported.

| | 5kb bulk GM12878 | | 5kb Micro-C H1-hESC | | 1kb Micro-C H1-hESC |
|---|---|---|---|---|---|
| | Unfiltered | Filtered | Unfiltered | Filtered | Unfiltered |
| JOnTADS | 37268 | 1774 (21 chromosomes) | 10304 | 1311 (18 chromosomes) | 4228 (chromosome 22) |
| Stripenn | 80121 | 1981 | 40734 | 5563 | NA |

**Supplementary Table 7**: The numbers of unfiltered and filtered stripes for 5kb bulk GM12878, 5kb Micro-C H1-hESC, and 1kb Micro-C H1-hESC data. Filtered: stripes passing the Stripenn p-value threshold of $< 0.1$; Unfiltered: all identified stripes without filtering. When calculating Stripenn p-values for filtering stripes identified by JOnTADS, Stripenn encounters errors for certain chromosomes. Therefore, the number of filtered stripes is reported only for chromosomes where Stripenn can be successfully applied. Additionally, since Stripenn does not support data finer than 5Kb resolution, we could not call stripes from Stripenn for 1kb Micro-C hESC data and Stripenn p-values for filtering stripes identified by JOnTADS could not be obtained. Thus, only unfiltered striped identified by JOnTADS were reported for 1kb Micro-C H1-hESC data.
